# USP15 deubiquitinase safeguards hematopoiesis and genome integrity in hematopoietic stem cells and leukemia cells

**DOI:** 10.1101/2020.01.23.916627

**Authors:** Paul van den Berk, Cesare Lancini, Carlos Company, Michela Serresi, Danielle Hulsman, Colin Pritchard, Ji-Ying Song, Matthias Jürgen Schmitt, Ellen Tanger, Ivo J. Huijbers, Heinz Jacobs, Maarten van Lohuizen, Gaetano Gargiulo, Elisabetta Citterio

## Abstract

Altering ubiquitination by disruption of individual deubiquitinating enzymes (DUBs) has proven to affect hematopoietic stem cell (HSC) maintenance. However, comprehensive knowledge of DUB function during hematopoiesis *in vivo* is lacking. To accomplish this goal, we systematically inactivated DUBs in mouse hematopoietic progenitors using *in vivo* small hairpin RNAs (shRNAs) screens. We found that multiple DUBs may be individually required for hematopoiesis and that the ubiquitin-specific protease 15 (USP15) is particularly important for the maintenance of murine hematopoietic stem and progenitor cells *in vitro* and *in vivo.* Consistently, *Usp15* knockout mice exhibited a reduced HSC pool. The defect was intrinsic to HSCs, as demonstrated by competitive repopulation assays. Importantly, USP15 is highly expressed in normal human hematopoietic cells and leukemias, and USP15 depletion in murine early progenitors and myeloid leukemia cells impaired *in vitro* expansion and increased genotoxic stress. Our study underscores the importance of DUBs in preserving normal hematopoiesis and uncovers USP15 as a critical DUB in safeguarding genome integrity in HSC and in leukemia cells.

## Introduction

Mammalian hematopoiesis depends on a limited pool of stem cells in the bone marrow. Hematopoietic stem cells (HSCs) have the unique properties of self-renewal and multi-lineage potential, giving rise to daughter stem cells and to committed progenitors, thereby achieving lifelong hematopoiesis. This is accomplished by careful maintenance of a homeostatic balance between HSC quiescence, self-renewal and differentiation (de Haan and Lazare, 2018; Laurenti and Gottgens, 2018; Morrison and Spradling, 2008). Perturbation of this balance as well as replication stress can cause stem cell failure or transform normal HSCs and progenitors into disease-initiating leukemic stem cells (LSCs) (Flach et al., 2014; Warr et al., 2011). Understanding HSC and bone marrow homeostasis is therefore essential for understanding mechanisms controlling diseases and ultimately for targeting LSCs (Warr et al., 2011).

The 76 amino-acid molecule ubiquitin is conjugated to proteins as a monomer (mono-ubiquitination) or in the form of ubiquitin-chains (poly-ubiquitination) through the sequential action of E1 ubiquitin activating-, E2 ubiquitin conjugating- and E3 ubiquitin ligating-enzymes (Yau and Rape, 2016). Deubiquitinating enzymes (DUBs; also referred to as deubiquitylating enzymes or deubiquitinases) reverse substrate ubiquitination. By removing or editing the ubiquitin chains from their targets, DUBs critically regulate ubiquitin-mediated signaling pathways, including protein homeostasis, DNA repair and receptor signaling (Clague et al., 2012; Mevissen and Komander, 2017). Consequently, deregulation of DUBs is implicated in human pathologies, such as cancer, neurodegenerative and hematological and infectious diseases (Heideker and Wertz, 2015).

The human genome encodes ≈100 DUBs, grouped in seven families based on structural properties. Four of these DUB families belong to the cysteine proteases [ubiquitin-specific proteases (USPs), ovarian tumor proteases (OTUs), ubiquitin C-terminal hydrolases (UCHs) and Josephin domain DUBs], one is the motif interacting with ubiquitin (MIU)-containing novel DUB family [MINDYs], one consists of the JAMM/MPN+ family of metalloproteases, and the seventh is represented by the recently identified ZUFSP (Haahr et al., 2018; Kwasna et al., 2018; Mevissen and Komander, 2017). By a mouse genetic approach our laboratory reported that loss of the ubiquitin-specific protease 3 (USP3) protects the HSCs from functional decline through modulation of ubiquitin-dependent DNA damage response (DDR), a critical genome maintenance pathway (Lancini et al., 2014). This is in line with a proper DDR being crucial to HSC function (Bakker and Passegue, 2013; Biechonski et al., 2017). Notably, numerous DUBs control ubiquitin-dependent DDR signaling (Citterio, 2015; Jackson and Durocher, 2013; Nishi et al., 2014; Schwertman et al., 2016) and recent studies provided evidence that DUBs deregulation contributes to altered HSC homeostasis and human blood diseases, including myelodysplastic syndrome (MDS) (Adorno et al., 2013; Dey et al., 2012; Gu et al., 2016; Higuchi et al., 2016; Nakagawa et al., 2015), with some of the DUBs primarily impacting on DDR (Citterio, 2015; Lancini et al., 2014; Parmar et al., 2010).

Functional analysis of HSC within their physiological environment is more likely to result in finding modulators potentially relevant in disease (Morrison and Spradling, 2008; Schepers et al., 2015). Unbiased, functional genomic approaches by short hairpin RNAs (shRNAs) have demonstrated the power of forward RNAi screens in dissecting functional aspects of both normal (Cellot et al., 2013; Galeev et al., 2016; Hope et al., 2010) and leukemic HSCs (Zuber et al., 2011). Using lentiviral-based libraries frequently employed in cancer gene discovery (Gargiulo et al., 2014), pooled *in vivo* screening approaches in early murine hematopoietic precursors led to the identification of critical factors limiting normal HSC self-renewal under DNA damage conditions (Sertorio et al., 2017; Wang et al., 2012) as well as of determinants of malignant hematopoiesis (Miller et al., 2013; Puram et al., 2016).

While recent gene-centric approaches connected DUBs to HSC maintenance (Adorno et al., 2013; Dey et al., 2012; Gu et al., 2016; Higuchi et al., 2016; Lancini et al., 2014; Nakagawa et al., 2015; Nijnik et al., 2012; Parmar et al., 2010; Wang et al., 2013), a comprehensive understanding of DUBs biological functions in hematopoiesis and leukemia is missing. DUBs are poorly represented in *in vivo* published screens (Sertorio et al., 2017; Wang et al., 2012) and *in vitro* functional approaches for DUBs in cancer cell lines were hypothesis driven (Nishi et al., 2014). In this study, we individually depleted all DUB genes using *in vivo* RNAi screens in mouse hematopoietic precursors, with the aim of ranking the most relevant DUBs required for normal and malignant hematopoiesis. From this systematic and targeted approach, we uncovered novel and known DUBs as putative regulators of hematopoietic precursors activity and highlighted USP15 as a novel determinant of hematopoiesis *in vivo* and its role in preserving genome integrity, with potential implications for combinatorial treatments in leukaemia.

## Results

### *In vivo* RNAi screens for deubiquitinating enzymes (DUBs) identify known and novel regulators of hematopoietic stem and progenitor cell activity

To identify DUB determinants of mouse HSC activity, we performed pooled *in vivo* RNAi screens using adult murine hematopoietic stem and progenitor cells in a bone marrow (BM) transplantation setting (Fig. 1A). To this end, we generated a custom pool of 508 lentiviral shRNAs vectors potentially targeting all annotated mouse orthologues of human DUBs (≈100) (Mevissen and Komander, 2017). This primary library contained 3 to 6 shRNA vectors per gene, selected from the shRNA library developed by the RNAi Consortium (TRC) at the Broad Institute (Open Biosystem) (Table S1, S2). Since the statistical representation of shRNA libraries is critical for success in *in vivo* screening, we used the full library in a primary screen, and divided the library in two sub-pools (DUB1 and DUB2 sub-libraries), subsequently used in secondary screens (Fig. 1B). To perform qualitative controls, we included in each library shRNAs targeting known HSCs regulators as positive controls (Park et al., 2003; Vasanthakumar et al., 2016; Wang et al., 2012).

**Figure 1.**
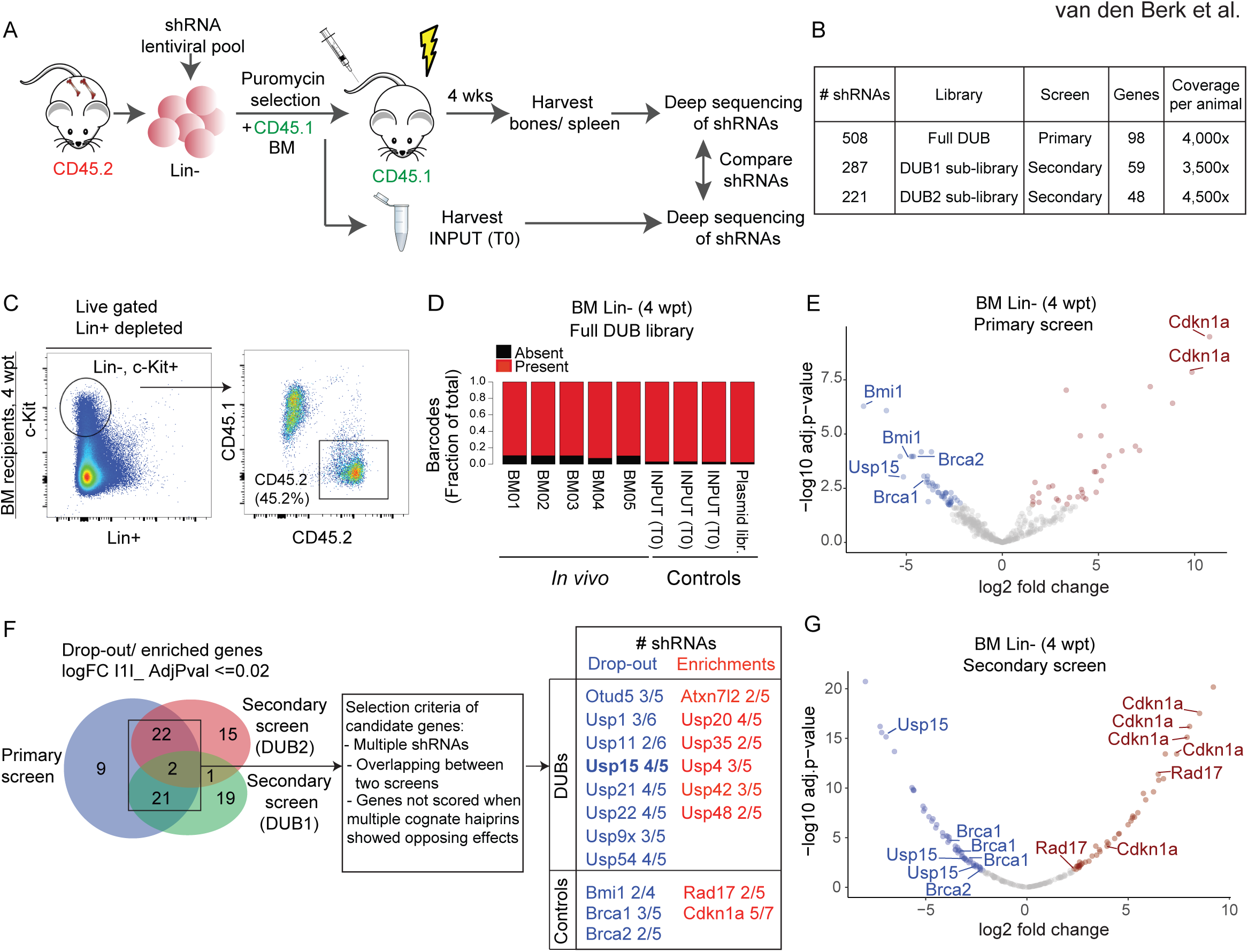
Pooled *in vivo* RNAi screen identifies candidate DUBs effectors of hematopoietic stem and progenitors cell activity. (A) Overview of the DUB RNAi screens *in vivo*. Lineage negative (Lin-) cells isolated form wt (CD45.2) bone marrow (BM) were grown under stem cell conditions and transduced with pooled lentiviral libraries targeting all annotated mouse orthologues of human DUBs (≈100). After selection, CD45.2 transduced Lin-cells are mixed 1:1 with CD45.1 support BM cells. A fraction of the transduced CD45.2 cells, INPUT (T0), is isolated before injection. Proviral shRNAs inserts were sequenced from BM isolated Lin-cells and splenocytes from recipient mice at 4 weeks post transplantation (wpt) and from INPUT cells to determine the changes in distribution of all shRNAs. (B) The primary screen was performed with the full shRNA DUB library in five replicate mice. The DUB library was divided in two sub-pools, DUB1 and DUB2 sub-libraries, and two secondary screens were performed. Most of the targeted genes do not overlap between DUB1 and DUB2 sub-libraries. (C) shRNA library-transduced Lin-cells efficiently engraft and contribute to BM repopulation. The Lin-fraction was purified from recipient mice at 4 wpt and analyzed for chimerism. Representative FACS profiles are shown. (D) The primary screen shRNAs library complexity is maintained *in vivo*. Consolidated fraction of shRNAs retrieved *in vivo* in Lin-cells at 4 wpt and in controls. (E) Vulcano plot depicting the log2 fold change in the BM of recipient mice of all hairpins used in the primary screen, normalized across five replicates. (F) Venn diagram depicting significantly differentially represented genes overlapping between the primary and secondary screens. Candidate genes were selected with the indicated criteria. (G) Vulcano plot depicting the log2 fold change in the BM of recipient mice of all hairpins used in the secondary screen (DUB2 sub-library), normalized across seven replicates. (E-G) Significantly (AdjPval ≤0.02) dropout (log2 FC≤ 1, blue) and enriched (log2FC ≥1, red) shRNAs are highlighted. See also Figure S1 and S2.

Freshly isolated Lineage negative (Lin-) BM cells were transduced with the titered shRNAs pooled library (MOI<<1), selected with puromycin and subsequently injected into lethally irradiated mice (Fig. 1A). In this limited time window, Lin-cells were maintained *in vitro* in the presence of HSC cytokines, in conditions that are known to preserve and enrich for stem cells/early progenitors (Materials and Methods)(Ye et al., 2008). Indeed, early progenitors were maintained during transduction, as gauged by the enrichment of the Lin-cKit+Sca1+ (LSK) cells in FACS analysis (Fig. S1A). Notably, the transduced cell culture also retained phenotypic HSCs, which was assessed by the HSC SLAM surface marker CD150+ that is expressed on cells endowed with an immature phenotype and reconstitution potential (Christensen and Weissman, 2001; Kiel et al., 2005; Yeung and Wai Eric So, 2009) (Fig. S1A). Transduced Lin-cells were mixed 1:1 with total BM cells from CD45.1 mice, which is the state of art for competitive *in vivo* transplantation (Fig. 1A). To ensure optimal representation of the shRNA library, we have injected a minimum of 1×10^6^ Lin-transduced cells/mouse, aiming at least at a predicted 2000-fold library representation per animal. This representation is estimated to be sufficient to control for grafting efficiency and stochastic drifts (Gargiulo et al., 2014).

We allowed cells to engraft recipient animals and harvested blood, BM and spleen from recipient mice at 4 weeks post transplantation (wpt). We chose a 4-week time-point as read-out of the screen based on experimentally determined parameters. First, we verified that 4 weeks is a sufficiently long time to allow the assessment of potential phenotypic defects of the murine progenitors during the acute proliferative phase. This included both expansion and depletion, thereby enabling us to identify genes regulating either quiescence or proliferation. Second, 4 weeks is a time-frame consistent with polyclonal engraftment and insufficient to allow manifestation of compensatory mechanisms and HSC clonality issues. In fact, in long-term engraftment experiments (4-6 months), only a small number of HSC contribute to most cellular output (Naik et al., 2013). In our experiments, we observed measurable grafting in recipients and the generation of donor-derived B cells in the spleen of transplanted recipients (Fig. S1). This supports the 4-weeks time-point as being sufficient to enable the screen while limiting HSC clonal expansion.

FACS analysis of BM, circulating blood cells and splenocytes showed successful engraftment of the transduced Lin-cells, with an average of 50% contribution as measured in the BM (Fig. 1C, Fig. S1B and S1C). To assess the relative representation of each shRNA *in vivo*, we then performed parallel next generation sequencing of PCR-amplified shRNA sequences from genomic DNA in the following conditions: 1) *in vivo* hematopoietic precursors and differentiated cells, isolated at 4 wpt from the BM (Lin-cells) or the spleen (CD43-, CD45.2+, CD19+, CD220+ B cells), respectively, of recipients, 2) controls: transduced Lin-cells immediately before injection (INPUT, T0) as well as the plasmid library. Sequencing of individual samples revealed that individual shRNA abundance in transduced Lin-(T0) correlated well with the hairpin reads in the plasmid library, supporting efficient transduction *in vitro* (R2=0.69, Fig. S2A). Importantly, more than 97% of the hairpins could be identified in the transduced Lin-(T0) and more than 89% were retrieved *in vivo* in purified Lin-cells from each recipient mouse (4 wpt). We concluded that a significant proportion of the initial library complexity is maintained *in vitro* and *in vivo*, respectively (Fig. S2A and Fig. 1D).

Principal component analysis (PCA) showed that the five *in vivo* BM samples were more similar to each other and were distinct from the input cells before injection, and limited variance between the individual samples was found (Fig. S2B, C). Moreover, a positive correlation was found between the relative representation of shRNAs retrieved from the BM to the ones retrieved from the spleen (R2=0.668), thereby supporting an overall good quality of the primary screen (Fig. S2D and Table S3).

Next, we performed a differential enrichment analyses on the *in vivo* and control samples. Overall, the behavior of well-known HSC regulators enhanced confidence in our *in vivo* RNAi screen. In fact, among the top hits we found genes relevant to HSC biology to be either enriched (involved in cell cycle restriction) or depleted (supporting self-renewal), including our positive controls. Consistent with the requirement for Bmi1 in adult HSC self-renewal (Park et al., 2003), two out of the 4 shRNAs targeting Bmi1 showed significant dropout (> 20 fold) in Lin-cells *in vivo* (Fig. 1E, Table S3). DNA repair genes BRCA1 and BRCA2/FANCD1 were also highly depleted with at least one shRNA per gene, in line with their crucial role in HSC survival (Navarro et al., 2006; Vasanthakumar et al., 2016). Consistent with a role in cell cycle restriction (Wang et al., 2012), two shRNAs for the cell cycle inhibitor *Cdkn1a* were enriched (Fig. 1E). Notably, DUBs with established importance in HSC maintenance, including USP1 (Parmar et al., 2010), USP3 (Lancini et al., 2014) and USP16 (Adorno et al., 2013; Gu et al., 2016) also scored top hits from the primary screen, and were targeted by two independent shRNAs (Fig. 1F, Fig. S2H and Table S3). These data further demonstrate the ability of our unbiased *in vivo* screen to identify DUBs relevant to HSC biology.

To validate our primary screen and increase its statistical power, we divided the primary library in two, mostly non-overlapping, shRNA sub-pools (DUB1 and DUB2 sub-library) and performed secondary screens under similar transplantation conditions (Fig. 1B, 1F and Fig. S1C). In line with the primary screen, high hairpins representation *in vitro* and *in vivo* (>95%), low variance between individual mice and the performance of positive control shRNAs support the overall quality and reproducibility of the secondary screens as well (Fig. 1F, G, Fig. S2E-G, and Table S3). Although many shRNAs showed similar changes in representation in the primary and in the secondary screens, a measurable variation was also present. This is likely due to inconsistencies in transduction efficiency or to the stochastic gain or loss of shRNAs following *in vivo* growth (Table S3). To overcome this, we adopted stringent selection parameters. We considered as candidates for follow up analyses those genes for which at least two shRNAs were depleted/enriched by 10-fold median in the BM relative to their representation in the T0 control (i.e. the injected cell population; Adj.Pvalue ≤0.02) in each screen, and which were called as hits in at least two independent experiments. When multiple hairpins showed opposite effect, the corresponding gene was excluded. By these criteria, our positive controls as well as 14 out of 81 DUB genes tested were validated in the secondary screens and were defined as positive hits (Fig. 1F, Fig. S2H).

To prioritize hits for follow up, we next focused on DUBs with reported high expression in LSK and in HSC (Cabezas-Wallscheid et al., 2014; Heng et al., 2008; Lancini et al., 2016) (immgen.org). We choose to focus on USP15, for which three independent shRNAs were depleted for >15-fold median in the bone marrow after 4 weeks, and the top scoring shRNAs showed a 60-fold dropout (Fig. 1E and G, Table S3). We and others found that USP15 (Baker et al., 1999) is expressed in the early progenitor compartment (LSK) and in HSC (Cabezas-Wallscheid et al., 2014; Lancini et al., 2016), as well as in blood and in splenic B cells (Lancini et al., 2016; Vlasschaert et al., 2017). Together with our screen results, these data suggest a potential role for USP15 in hematopoiesis, though no functional study *in vivo* has yet been reported. We therefore decided to further investigate the role of USP15 in HSC biology.

### USP15 depletion impairs hematopoietic stem and progenitor cells proliferation *in vitro*

To investigate whether a role for USP15 in hematopoietic progenitors self-renewal is plausible, we first checked USP15 expression levels in the normal hematopoietic developmental tree by surveying published gene expression data sets. In the mouse bone marrow, *Usp15* expression is consistently high at single cell level and expression is homogeneous in the entire hematopoietic tree, being expressed at similar level in single mouse LT-HSCs and early lineage-committed progenitors (Fig. S2I, J) (Nestorowa et al., 2016; Olsson et al., 2016). Importantly, *Usp15* expression pattern in the mouse is similarly conserved in humans, as inferred by *USP15* expression in CD34+ human HSC as well as in early lineage-committed progenitors at single cell level (Fig. S2K) (Pellin et al., 2019).

We therefore addressed the impact of individual USP15-targeting shRNAs on hematopoietic progenitors *in vitro* and *in vivo* (Fig. 2A). We first assessed the ability of the single shRNAs to reduce *Usp15* expression upon low multiplicity of infection (MOI<<1), similar to the transduction conditions used in the RNAi screens. To cope with paucity of Lin-cells, we chose RT-qPCR as readout for knockdown efficiency. All three shRNAs identified in the secondary DUB screen (DUB2 sub-library; Fig. 1G, Table S3) downregulated USP15 expression in freshly isolated, lentiviral-infected Lin-cells (Fig. 2B). To functionally validate the two top-scoring lentiviral shRNA vectors in the screen, Lin-cells were transduced with either a control (shScramble) or USP15-targeting #sh16 and #sh17 shRNAs. To determine the effect of USP15 depletion on the LSK compartment, the transduced cells were propagated in a serum-free medium supplemented with pro-self-renewal growth factors and analyzed by flow cytometry for the presence of LSK surface receptors at one-week post infection (8 days from initial culture). Within the Lin-, c-Kit+ population, the fraction of LSKs, representing the undifferentiated compartment capable of *in vivo* repopulation, remained comparable between the USP15-depleted and the control shRNA cells (Fig. 2C, left panel). Nevertheless, the expansion of both Lin-, c-Kit+ and of LSK cells was affected by USP15 depletion compared to control shRNA (Fig. 2C, middle and right panels and Fig. S3A). Consistently, USP15 knockdown progenitors exhibited limited proliferation (Fig. 2D). Altogether, these data show that *in vitro* reduced expression of USP15 affects the proliferative ability of hematopoietic progenitors.

**Figure 2.**
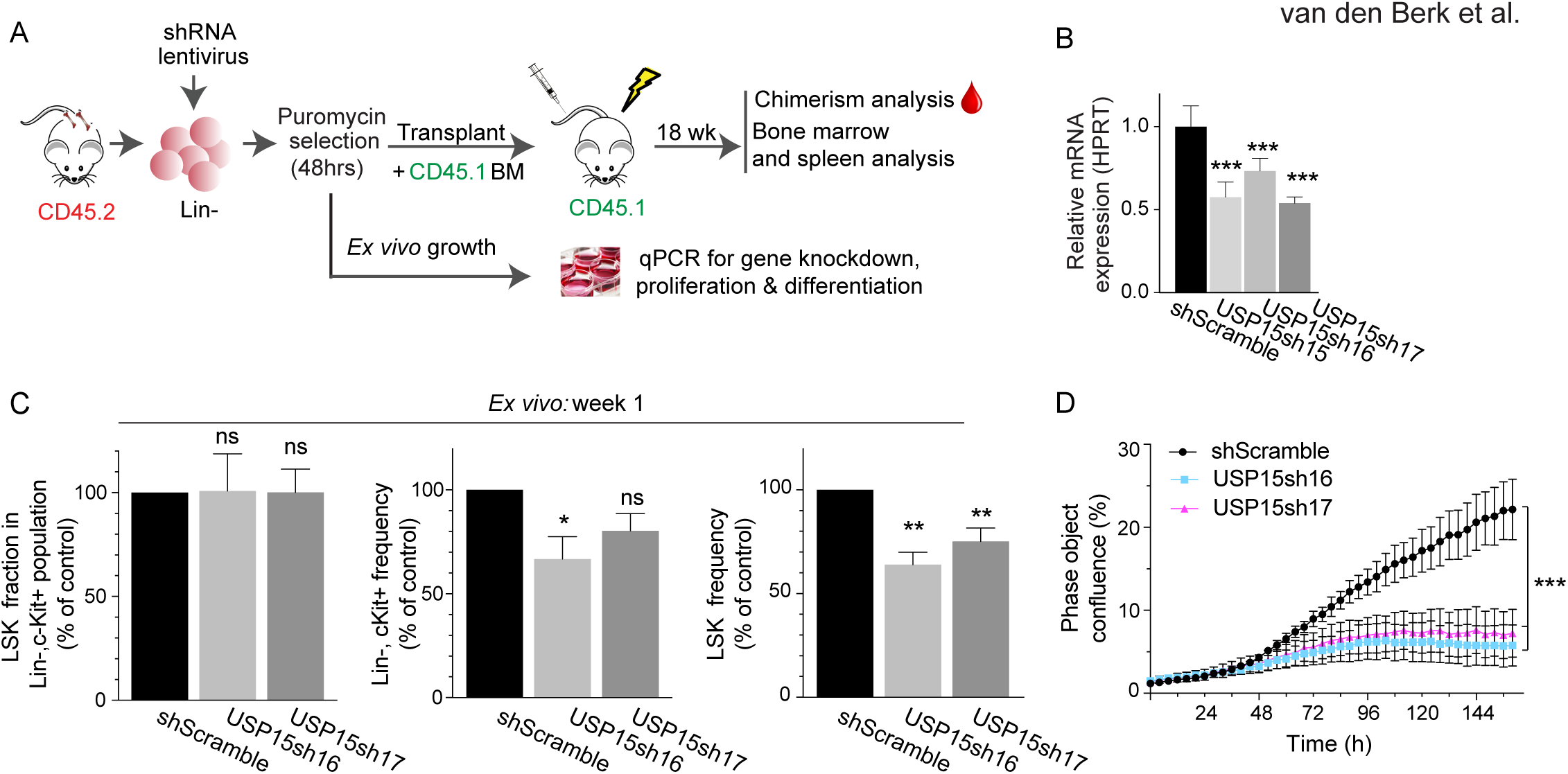
Knockdown of USP15 in hematopoietic stem and progenitor cells reduces their proliferation capacity *in vitro*. (A) Overview of the *in vitro* and *in vivo* validation assays for USP15-targeting shRNAs. (B) Freshly isolated Lin-cells were infected with non-targeting (shScramble) or USP15-targeting (shUSP15) shRNAs expressing lentiviruses. The knockdown efficiency of USP15 was detected by real-time qPCR using two different pairs of USP15 primers. HPRT was used as the internal control for normalization. Mean values of three technical replicates ± SD are shown. (C) Flow cytometry analysis of Lin-cells at 1 week post infection with the indicated shRNAs. The frequency of LSKs in the Lin-, c-Kit+ population (left panel), and the frequency of Lin-, cKit+ and LSK in the live culture (middle and right panel, respectively) was calculated and normalized relative to shScramble control. N= 3 independent experiments. Mean values ± SEM are shown. FACS representative profiles are presented in Figure S4. (D) Freshly purified Lin-cells were plated 7 days post infection with the indicated shRNAs and monitored for growth. Kinetic measures the number of cells, recorded over time and plotted as phase contrast object confluence. *n* = 4 wells per data point. Mean values ± SEM are shown. For all panels: *, P ≤ 0.05; **, P ≤ 0.01; ***, P ≤ 0.001. P-value was assessed by Student’s *t* test (A-C) or Multiple t test (D) in Prism 7. See also Figure S3A.

### USP15 depletion impairs stem and progenitor cells reconstitution potential *in vivo*

To assess the effect of USP15 knockdown in a physiological setting, we transduced murine Lin-progenitors with USP15-targeting or control shRNAs and competitively co-transplanted these CD45.2 USP15-depleted or control progenitors together with freshly isolated CD45.1 BM cells (1:1 ratio) into lethally irradiated recipients. Chimerism in the blood was monitored over time. Within a period of 18 weeks, USP15 knockdown Lin-cells failed to contribute to a chimerism level beyond the 20% of total peripheral blood cells, whereas the chimerism level of control mice progressively increased, reaching the expected approximate 50% contribution (Fig. 3A). This underscores a competitive disadvantage of USP15 depleted cells compared to control shRNA-transduced cells. At 18 wpt, we found that all lineages within CD45.2 USP15-depleted peripheral blood cells, including myeloid/granulocytes (CD11b+, GR1+ cells), B and T cells were equally affected as compared to their control counterpart (Fig. 3B and Fig. S3B). Similarly, USP15 loss affected multi-lineage reconstitution (B and T cells) of recipient animals’ spleen at 18 wpt, with an average 52% of control B cells compared to 25% and to 10.8% of USP15-#sh16 and USP15-#sh17 cells respectively (Fig. 3C and Fig. S3C).

**Figure 3.**
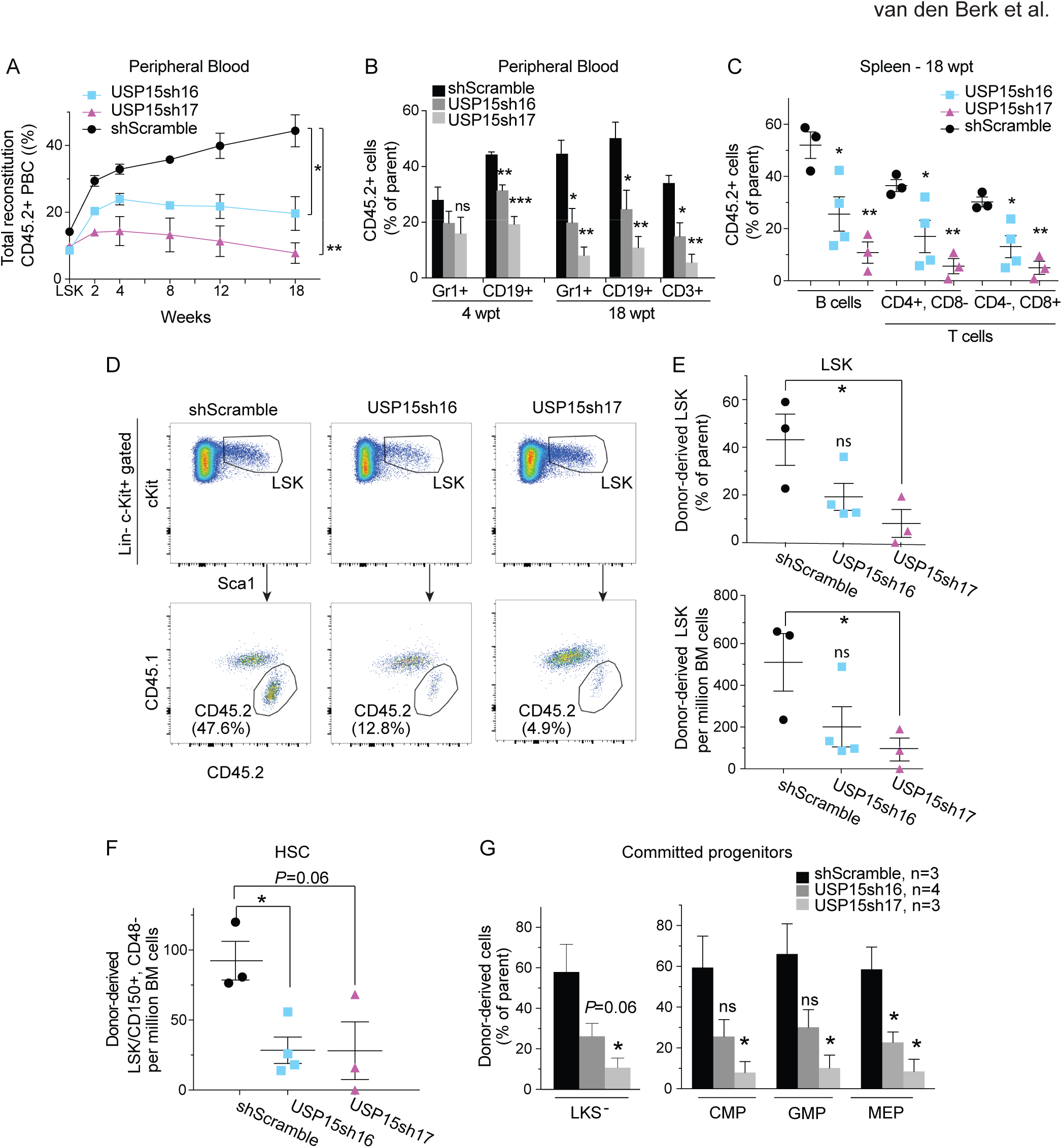
USP15 is required for HSC multilineage reconstitution potential into primary recipients. Freshly isolated wt murine Lin-cells were transduced with shRNAs targeting USP15 or control shRNA and assayed in competitive bone marrow transplantation. (A, B) CD45.2 chimerism in peripheral blood (A) and contribution of transduced cells to myeloid (Gr1^+^), B cell (CD19^+^), and T cell (CD3^+^) lineages in the blood (B) of recipients at the indicated time points. PBC, peripheral blood cells. (C) CD45.2 chimerism was measured in B and T cell lineages in the spleen of recipient mice at 18 wpt. (D) Representative FACS profiles of the Lin-, c-Kit+, Sca1+ (LSK) compartment and of chimerism analysis of this population in primary recipient mice at 18 wpt. (E) CD45.2 chimerism levels in LSK cells in primary recipients (upper panel). Lower panel shows the numbers of donor-derived LSK in 10^6^ viable BM cells at 18 wpt. (F) Cell numbers of donor-derived HSC (LSK/CD150+/CD48-) in 10^6^ viable BM cells at 18 wpt. (G) Fraction of donor derived progenitors populations (LKS^-^, CMP, GMP and MEP) in primary recipients at 18 wpt. Gating strategies and representative FACS profiles are presented in Figure S3. For all panels: *, P ≤ 0.05; **, P ≤ 0.01. P-value was assessed by Student’s *t* test in Prism 7. Data represent mean values ± SEM. N=3 per shRNA, except for shUSP15#16, n=4.

The above results suggest a defect in the multilineage reconstitution potential of USP15 depleted progenitors. Given that BM resident HSCs are mainly responsible to give rise to and maintain all blood cell lineages (Kiel et al., 2005; Wilson et al., 2008), we next quantified the numbers of CD45.2+ cells in the BM of recipient mice transplanted with either USP15-depleted or control progenitors (Fig. S3D). To assess stem cell reconstitution, we chose 18 wpt as time-point for BM immuno-phenotypical analysis, a time which is sufficient for essentially all mature hematopoietic cells to be derived from the most undifferentiated HSC (Naik et al., 2013). In line with the overall lower relative contribution to blood reconstitution (Fig. 3A), we measured a defect in USP15-depleted BM precursors. In fact, USP15-depleted LSK were reduced in frequency and numbers (2.38 and 8-fold reduction, respectively) compared to control (shScramble) LSK, which reached 50% contribution to the recipient mice LSK compartment (Fig. 3D, E). To specifically focus on HSCs, we then employed the HSC surface receptors SLAM CD48 and CD150 markers (Cabezas-Wallscheid et al., 2014; Kiel et al., 2005; Oguro et al., 2013). We found a significant decline (3.25 fold) of CD42.2 HSCs (as defined by LSK/CD48-/CD150+) in the BM of animals reconstituted with USP15 depleted cells, compared to controls (Fig. 3F). These comparisons underscore a defect in long-term hematopoiesis caused by USP15 loss and a critical role for USP15 in HSC *in vivo* maintenance.

We next investigated the more differentiated, proliferative LKS^-^ progenitors. USP15 depletion resulted in a consistent decrease in donor-derived cells also in this compartment. A similar reduction of USP15-depleted cells compared to controls was measured in both the myeloid subsets of common myeloid progenitors (CMPs) and granulocyte-monocyte progenitors (GMPs) and in the megakaryocyte-erythrocyte progenitors (MEPs) (Fig. 3G and Fig. S3D) (Yeung and Wai Eric So, 2009), confirming an important role for USP15 in preserving all the main hematopoietic differentiation pathways.

In summary, *in vivo* phenotypic profiling supports a role for USP15 in preserving hematopoietic progenitor engraftment and multi-lineage reconstitution potential, which is needed for peripheral blood repopulation and distal hematopoiesis.

### USP15 knockout compromises normal HSC function *in vivo*

To assess the role of USP15 in physiological hematopoiesis, we generated mice deficient for USP15 by germline genetic ablation using the CRISPR-*Cas9* technology (Pritchard et al., 2017) (Fig. S4A). Deletion of the *Usp15* locus was achieved by co-injection of gRNAs and Cas9 mRNA into the pronuclei of mouse zygotes and confirmed by PCR genotyping and Western blot (Fig. S4B, C). Homozygous USP15 knockout mice were viable, indicating that USP15 is dispensable for embryonic development. However, initial characterization revealed that *Usp15-/-* animals were born at sub-Mendelian ratio and showed reduced survival when compared to *Usp15+/+* mice, confirming a critical role for USP15 *in vivo* (Fig. S4D, E). Decreased life span in *Usp15-/-* animals compared to wt littermates is accompanied by lower body weight (Fig. S4F). Interestingly, some of the *Usp15* KO animals showed evidence of inflammatory lesions (Fig. S4G, H, and Table S7).

We next screened young adult *Usp15+/+* and *Usp15-/-* littermates (8-14 wks) for BM cellularity. No marked differences were found, suggesting that USP15 deficient BM can develop to a large extent normally (Fig. S4I). In line with this, phenotypic analysis revealed a normal frequency in the Lin-, cKit+ population in *Usp15-/-* and control mice (Fig. 4A, D), with a modest (but not significant) reduction in the *Usp15-/-* more undifferentiated stem and progenitors, the LSKs (Fig. 4B, D). Notably, within LSKs, the frequency and numbers of immature precursors endowed with reconstitution potential (LSK, CD135-, CD150+) (Christensen and Weissman, 2001; Kiel et al., 2005; Yeung and Wai Eric So, 2009)(Fig. S4J-L) and, more specifically, of phenotypic HSC (LSK, CD48-, CD150+) (Cabezas-Wallscheid et al., 2014; Kiel et al., 2005; Oguro et al., 2013) (Fig. 4C, D) were significantly lower in knockout mice, reaching only 60% of their aged-matched wt controls. The more committed (myeloid) progenitor pools did not show any measurable phenotype, indicating that this compartment can functionally cope with a reduction of HSC at young adult age (Fig. S4M). Consistently, *Usp15-/-* BM cells performed similar to wt BM when assayed *in vitro* in myeloid colony formation assays (CFU-C) (Fig. S4N). In summary, during physiological hematopoiesis, USP15 loss particularly impacted on the HSC compartment, indicating that the maintenance of the more primitive HSC in adult mice depends on USP15 *in vivo*.

**Figure 4.**
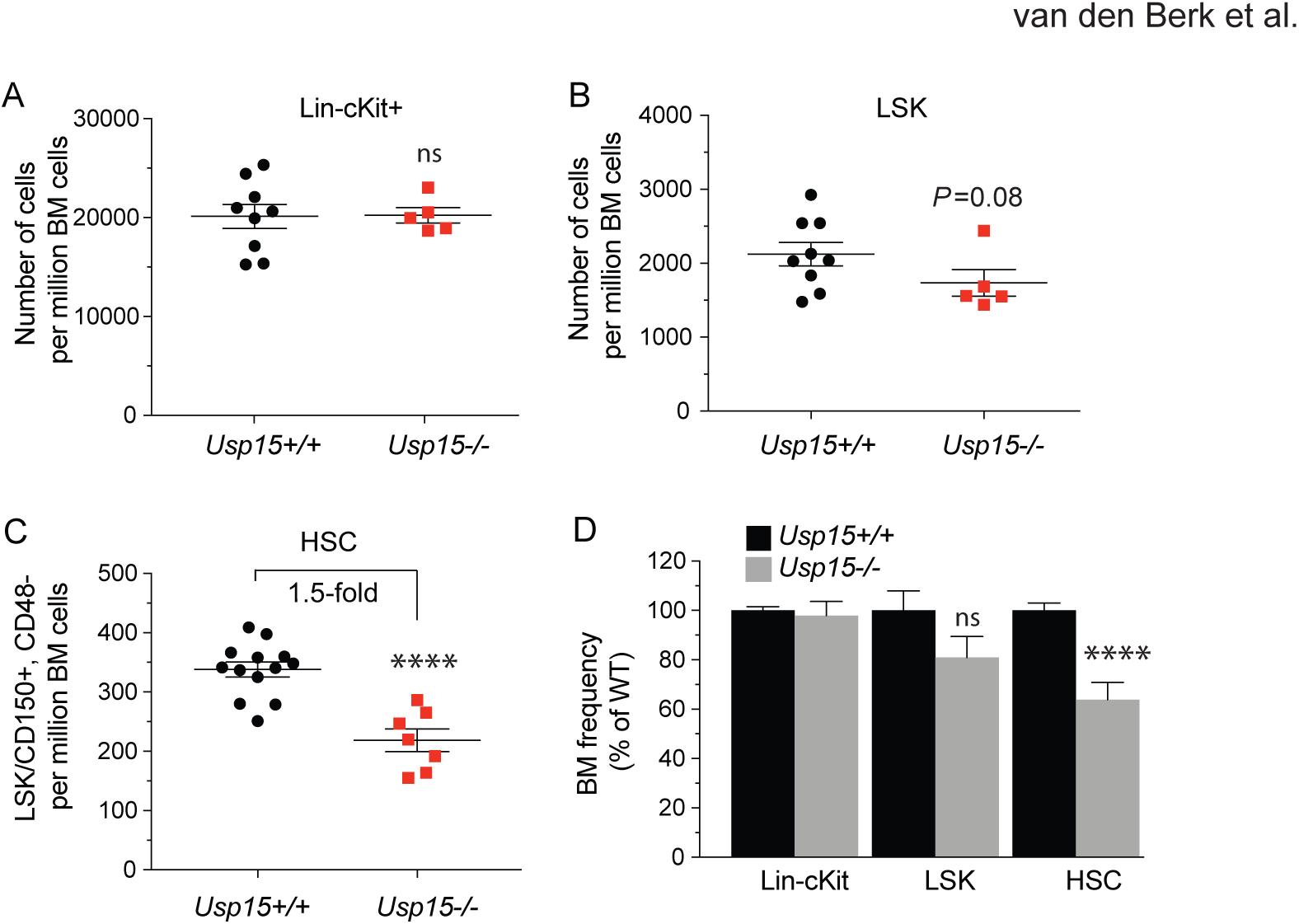
Reduced hematopoietic stem cell compartment in *Usp15* knockout mice. (A) Lin-, c-Kit+ cell numbers per million live BM cells. (B) LSK cell numbers per million live BM cells. (C) HSC (LSK, CD150+, CD48-) cell numbers per million live BM cells. (D) Frequency of Lin-, c-Kit+, LSK and HSC in bone marrow of *Usp15-/-* mice was calculated and normalized to *Usp15+/+* animals. Multiparameter flow cytometry analysis of the hematopoietic primitive populations was performed in 8-12 wks old *Usp15+/+* and *Usp15-/-* mice. Gating strategies are presented in Figure S3. Results are from three (Lin-, c-Kit+ and LSK; *Usp15+/+* n=9, *Usp15-/-n=5)* or four (HSC; *Usp15+/+* n=13, *Usp15-/-n=7*) independent experiments. For all panels: *, P ≤ 0.05, ****P ≤ 0.001 as assessed by Student’s *t* test. n.s.: not significant. Error bars represent ± SEM.

HSC are endowed with long-term multi-lineage reconstitution ability (Kiel et al., 2005). To establish whether the HSCs remaining in *Usp15* knockout mice are functionally equivalent to those in wt littermates, we performed competitive BM transplantations. Upon transplantation of BM cells containing a 1:1 mixture of test and competitor cells (Methods), chimerism of CD45.2 *Usp15-/-* peripheral blood cells in recipients significantly decreased over time compared to mice transplanted with *Usp15+/+* BM (Fig. 5A). *Usp15+/+* chimerism remained constant throughout the 18 weeks of analysis and reached the expected plateau. Importantly, USP15 deletion critically affected myeloid/granulocytes (CD11b+/Gr1+) as well as lymphoid blood cells (CD19+ B cells and CD3+T cells) (Fig. 5B). This phenotype recapitulates the USP15 knockdown defects observed upon transplantation of shRNA-transduced Lin-cells (Fig. 3A, B). In recipient BM at 18 wpt, a time shown to be sufficient for essentially all mature hematopoietic cells to be derived from the most undifferentiated HSC (Naik et al., 2013), we found significantly lower numbers of *Usp15-/-* LSKs as well as HSCs (LSK, CD150+, CD48-) compared to wt controls, suggesting that USP15-deficient HSCs have reduced self-renewal capacity into recipients compared to wt HSCs (Fig. 5C, D). Consequently, also the more committed *Usp15-/-* LKS^-^ and CMP pools were diminished (Fig. 5E).

**Figure 5.**
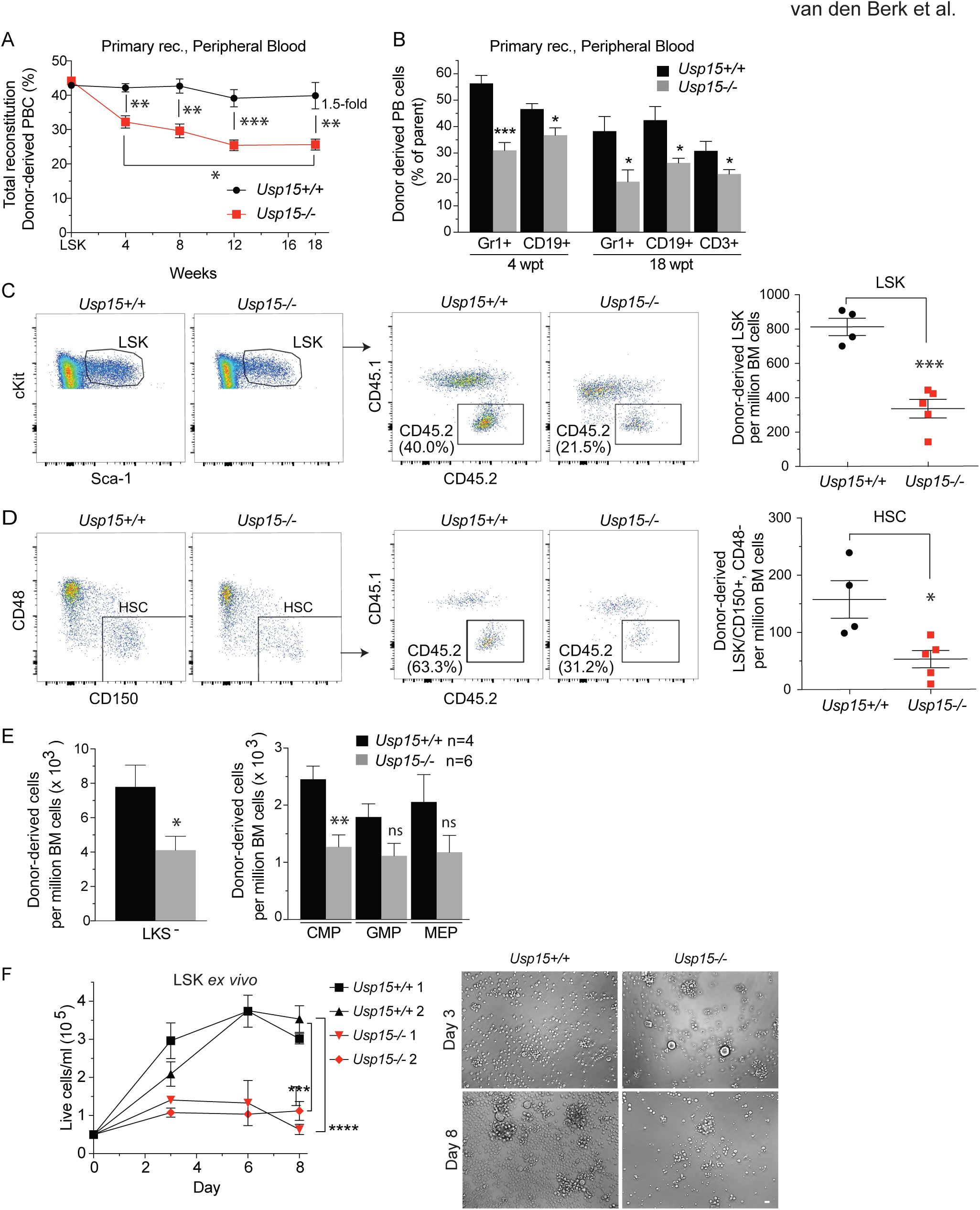
Genetic knockout of *Usp15* impairs HSC function. Competitive transplantation of BM cells freshly isolated from *Usp15+/+* or *Usp15-/-* mice. (A, B) Chimerism in peripheral blood (A) and contribution of BM cells to myeloid (Gr1^+^), B cell (CD19^+^) and T cell (CD3+) lineages in the blood (B) of recipients at the indicated time points. PBC, peripheral blood cells. (C) Representative FACS profiles (left panels) and numbers of donor-derived LSK per million viable BM cells in recipients at 18wpt. (D) Representative FACS profiles (left panels) and numbers of donor-derived HSC (LSK/CD150+/CD48-) per million viable BM cells in recipients at 18 wpt. (E) Numbers of donor-derived myeloid committed progenitor populations (LKS-, CMP, GMP and MEP) in recipients at 18 wpt. For all panels: *, P ≤ 0.05; **, P ≤ 0.01; ***, P ≤ 0.001. P-value was assessed by Student’s *t* test. Error bars represent ± SEM. (A-E): data represent one representative experiment out of 3. Recipient mice: (A, B): n=6 for each genotype; (C-E): *Usp15+/+* n=4, *Usp15-/-* n=5. (F) Freshly isolated, FACS sorted LSK were plated after 8d (second plating) in culture and monitored for growth. Cells were counted at the indicated days and the number of live cells is plotted. n = 4 wells per data point. Representative images at day 3 and 8 of the second plating are shown. Bar, 20 µm.

We next examined the consequences of USP15 deletion on HSPC cellular homeostasis. By DAPI/immuno-phenotyping combined analysis of freshly isolated BM cells, we measured that *Usp15-/-* mice have similar numbers of quiescent HSPCs compared to wt mice. The majority of HSCs were in the G0/G1 phase of the cell cycle. Under these physiological conditions, no subsets of the HSPCs nor the HSC differed significantly in terms of percentage of cells in S/G2 (Fig. S4O). Of note, freshly isolated *Usp15-/-* stem and progenitor cells did not show apparent apoptosis (Fig. S5P). Also, cleaved-caspase 3-positive cells were not readily detected on BM tissue sections of *Usp15-/-* mice (Fig. S4Q). RNA-seq of WT and *Usp15-/-* LSK confirmed the loss of *Usp15* and the maintenance of an overall stable identity of the cellular compartment (Fig. S4R).

Having established a functional defect in *Usp15-/-* LSK upon transplantation, we next assayed their intrinsic proliferative capacity in conditions of cytokine-induced replication. By monitoring the cell numbers of FACS sorted LSK in *in vitro* liquid cultures under stem cell conditions (Materials and Methods), we observed a significantly reduced intrinsic proliferative capacity for *Usp15-/-* LSKs compared to wt, which was exacerbated upon *ex vivo* culturing (Fig. 5F).

In all, systemic knockout validated USP15 as a novel positive regulator of HSC maintenance *in vivo* and proliferative capacity *ex vivo*.

### USP15 is highly expressed in human leukemia

Leukemic stem cells (LSCs) share functional properties with normal HSCs. For instance, acute Myeloid leukemia (AML) and Chronic Myeloid leukemia (CML) arise in the early hematopoietic compartment and have LSCs endowed with self-renewal and ability to propagate the disease (Kreso and Dick, 2014; Warr et al., 2011).

Consistent with this, USP15 featured the highest of expression in human hematopoietic tissues and related cancers, including leukemia and lymphomas (The Cancer Genome Atlas, TCGA)(Fig. 6A-B). In a AML-specific dataset, *USP15* expression was significantly higher in patients with AML carrying various genetic abnormalities compared to the normal human CD34+ -enriched BM hematopoietic precursors (Fig. 6C) (Bagger et al., 2013) (Hemaexplorer; www.bloodspot.eu). Of note, high expression of *USP15* is statistically associated with tissue-independent poor survival within the PANCANCER patient cohort, a feature generally associated with oncogenes (Fig. 6D and Table S8).

**Figure 6.**
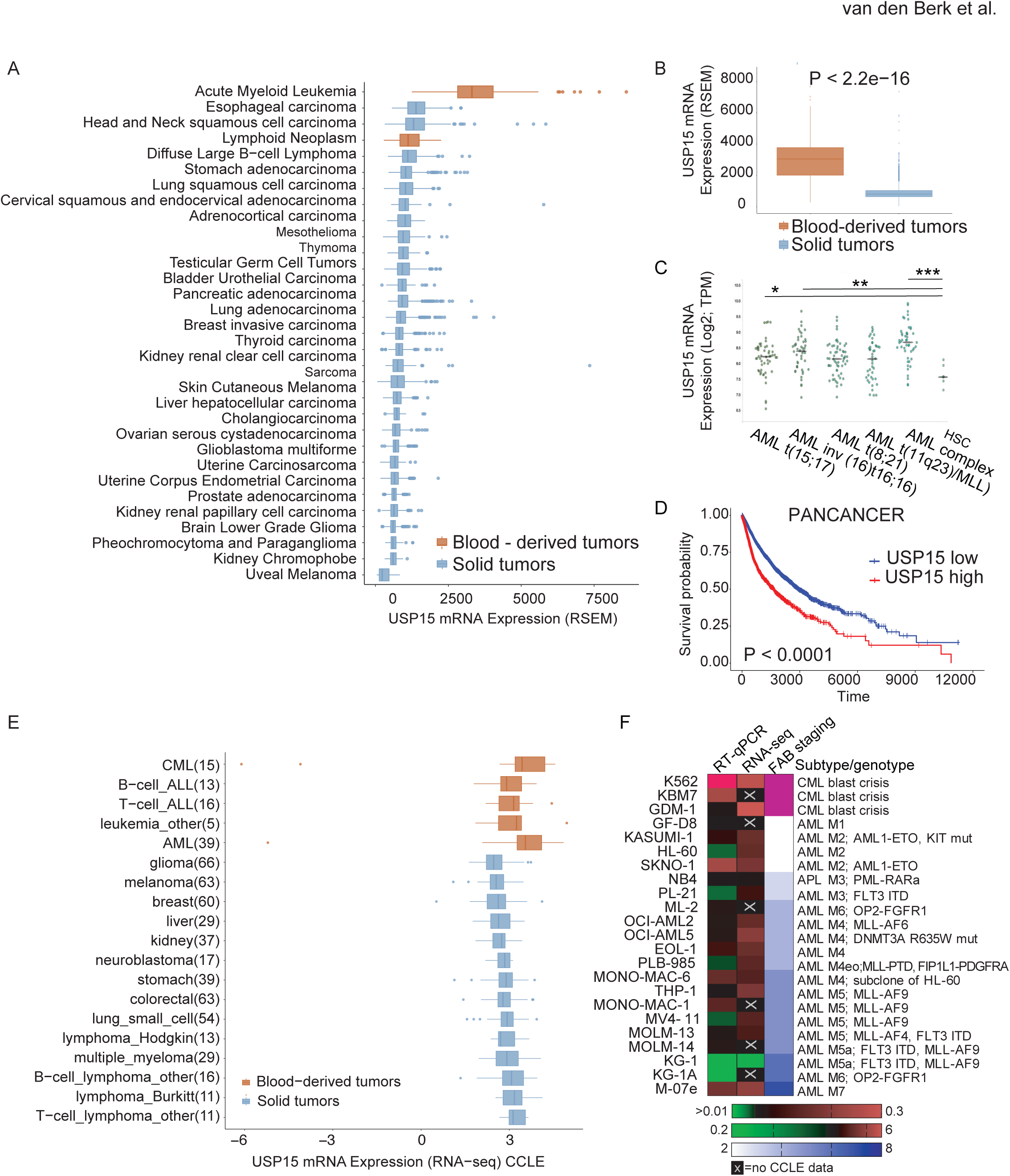
USP15 is highly expressed in primary blood-derived cancer. (A) Cohort TCGA Pan-Cancer (PANCAN)(total number of samples: 11060) shows up-regulation of *USP15* in Acute Myeloid Leukemia (AML). (B) Cohort TCGA PANCAN (11060) shows up-regulation of *USP15* in blood-derived tumors. (C) *USP15* is up-regulated in patients with AML. Log2 transformed expression of *USP15* from Microarray–Based Gene Expression Profiling of human BM cells (Hemaexplorer). TPM, Transcripts Per Kilobase Million. (D) High *USP15* levels are associated with poor survival. Kaplan-Meier curve correlating 10951 Pan-cancer patients’ survival with *USP15* gene expression is shown. TCGA Pan-Cancer (PANCAN) samples used in this analysis are shown in Table S8. (E) Expression of *USP15* from transcriptional profiling of human cancer cell lines (Cancer Cell Line Encyclopedia, Broad Institute) shows up-regulation of *USP15* in chronic myeloid leukemia (CML) and AML cell lines. (F) USP15 normalized expression levels in a panel of leukemia cell lines as assessed by RT-qPCR (left column) and RNA-seq by CCLE (middle column). The right column indicates the relative FAB stage for leukemia subtype classification.

To test whether these data are reflected in human cancer models, we next analyzed *USP15* expression in the large panel of comprehensively characterized Cancer Cell Line Encyclopedia (CCLE). In line with the previous analyses, the highest expression was found in leukemia cell lines, including multiple AML and CML cell lines, compared to all other tissues (Fig. 6E). To experimentally validate these analyses, we profiled USP15 expression in a twenty-three leukemia cell lines panel, including all maturation stages and chemotherapy resistant CML lines. With the sole exception of the KG1/KG1a cell line, USP15 mRNA was high in all the tested lines and independent from the leukemia stage. Interestingly, K562 and KBM7 blast crisis lines have very high USP15 expression (Fig. 6F).

To test whether USP15 gene expression correlates with its genetic dependency, we ranked the dependency scores calculated by Demeter2 for USP15 RNAi in CCLE lines (McFarland et al., 2018). According to DepMap (https://depmap.org), USP15 expression and dependency varied across cell lines but were not linearly correlated, nor leukemia cell lines were specifically sensitive compared to other cancers (Fig. S5A). Next, we investigated whether cancer-related biological pathways activation would be informative as a biomarker for USP15 dependency. To this end, we compiled a list of cell lines in which sensitivity to USP15 depletion was experimentally tested and could be classified as relatively high by DEMETER2 score (D2 score <-0.2) or low (D2 score >0.2). Among the leukemia cell lines, MV-4-11 and Kasumi-1 featured highly sensitive and SEM and K562 featured as less sensitive cell lines (Fig. S5A). Using PROGENy (Schubert et al., 2018), differential pathway activation between cell lines with varying degrees of sensitivity indicate that several RTK, JAK/STAT and PI3K signaling pathways tend to anti-correlate with sensitivity to USP15 depletion, whereas VEGFA, HIF1A and TGF-beta signaling were found more active in highly sensitive cell lines (Fig. S5B). Across the whole spectrum of CCLE cell lines, however, there was no evident biomarker for response, except a trend for activation of the Trail pathway (Fig. S5C), suggesting that USP15 depletion may operate in context dependent manner. To experimentally address the potential impact of the regulation of these pathways in response to USP15 depletion, we next performed RNAi of USP15 on highly expressing KBM7 and K562 CML cell lines. K562 are considered as a low sensitive cell line within the DepMap dataset and therefore response to USP15 RNAi may be uncoupled from survival. Ingenuity pathway analysis identified 657 and 330 differentially regulated genes in KBM7 and K562, respectively. In line with PROGENy analysis, RNAi of USP15 led to activation of inflammation-related pathways, which involve JAK/STAT and PI3K signal transduction (Fig. S5D-G). In K562, we also measured significant down-modulation of TGF-beta signaling (Fig. S5H, I), which is required for USP15 pro-oncogenic role in human glioma cells (Eichhorn et al., 2012).

Overall, these analyses indicate that USP15 is highly expressed in human primary leukemias and leukemia cell lines and that a subset of leukemia cells with various molecular alterations may be sensitive to USP15 loss.

### USP15 loss leads to genome instability in leukemia cell lines and in mice

Given the context-dependent responses to USP15 depletion in CML cells and that reversal of ubiquitination often contributes to fine-tuning of the DNA damage response (DDR) (Nishi et al., 2014), we next focused on exploring a potential role for USP15 in genome maintenance.

USP15 depletion by USP15-targeting siRNAs mildly but reproducibly reduced the viability of both “less sensitive” K562 and KBM7 and “more sensitive” MV411 and Kasumi-1 cell lines (Fig. 7A, B, S5A and below). Despite the predicted low sensitivity to USP15 depletion, USP15 loss was accompanied by significant increase in the number of spontaneous nuclear foci of the DNA damage response factor 53BP1 as well as increase in the basal levels of γ-H2AX DNA damage marker and in the frequency of micronuclei in both K562 and KBM7 (Fig. 7C-F), all indicative of enhanced genotoxic stress. This mirrors the increase in micronucleation, as well as bi- and multinucleation and apoptotic/necrotic cells observed in FACS sorted LSKs from the BM of *Usp15* knockout mice upon culturing (Fig. 7G), thereby indicating that USP15 loss affects genome integrity in all of these settings. Spontaneous genotoxic stress was also observed in USP15 depleted osteosarcoma cells (Fig. S6A-H), thereby extending the validity of USP15 expression as genome integrity safeguard mechanism to multiple tissue neoplasia.

**Figure 7.**
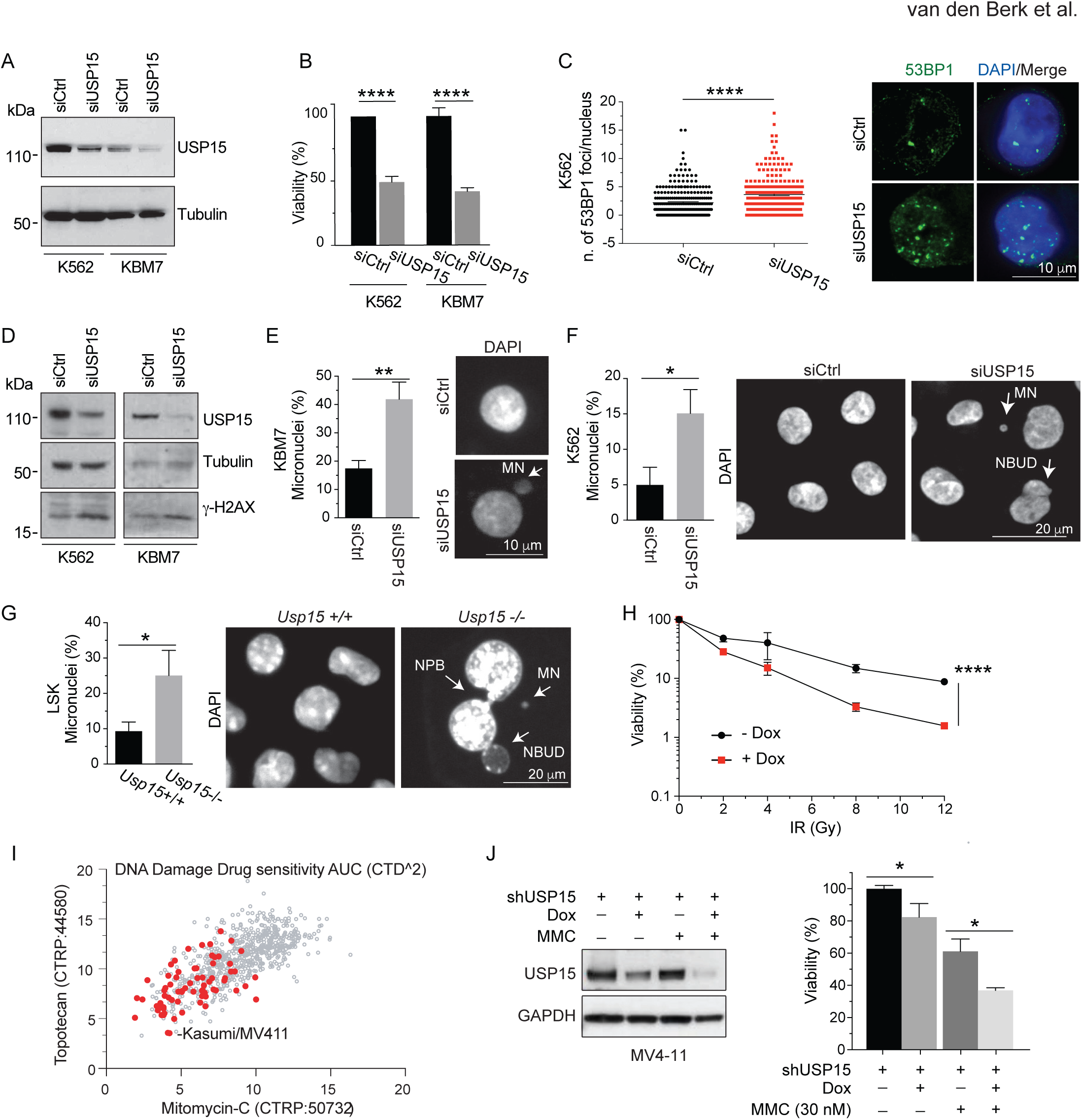
USP15 loss enhances genotoxic stress in human CML leukemic cells and in mouse normal hematopoietic progenitors. (A-F) K562 and KBM7 CML cell lines were transfected with USP15 (siUSP15) or non-targeting (siCtrl) siRNAs and assayed at 72hours after transfection. (A) Immunoblotting on whole cell extract with the indicated antibodies. (B) Viability in the indicated cell lines measured at 72 hours after transfection. Mean ± SD from three independent experiments is shown. (C) Representative images and quantification by Image J of the number of spontaneous 53BP1 foci per cell. Mean values ± SEM are shown. N=2. A minimum of 250 cells per sample was counted over two independent experiments. (D) Immunoblotting of USP15-depleted cell lines at 72 hours after transfection. Levels of γH2AX and tubulin were assayed. (E, F) USP15 depletion significantly increases micronuclei formation in KBM7 (E) and K562 (F) cell lines. Quantification and representative images of micronuclei (arrows) are shown. Results are mean ± SD from 3 independent experiments. A minimum of 150 (KBM7) or 450 (K562) cells was scored. (G) Percentage of micronuclei in FACS sorted, murine LSK after 11 d in culture. A minimum of 60 cells/genotype was scored in two independent experiments (each exp. *Usp15+/+* n=3; *Usp15-/-* n=2). Mean ± SD is shown. (H) KBM7 cells transduced with a doxycycline (dox)-inducible shUSP15 were kept in medium with or without dox for 5 days. Cells were then seeded for IR treatment with the indicated dose. Cell viability was measured by Alamar Blue 3d after IR. Mean ± SD of two independent experiments (each with n=5 replicates per sample) is shown, two-way analysis of variance (ANOVA) ****p <0.0001. Similar data were obtained with siUSP15. (I) Scatter plot of area under the dose–response curve (AUC) scores indicating sensitivity of individual cell lines to either Topotecan or Mitomycin-C. Red dots indicate leukemia cell lines. Data are generated by Cancer Target Discovery and Development (CTD2) Network and taken from the Cancer Therapeutics Response Portal (CTRP). (J) MV4-11 cells harboring USP15 shRNA were kept in medium with or without doxycycline (Dox) for 5 days and subsequently plated with the indicated concentration of Mitomycin-C (MMC). Western blot (left panel) and cells viability (right panel) were performed at 72 hours of MMC treatment. Results are the mean ± SEM of three (-MMC) or two (+MMC) independent experiments (each with n=3 replicates per sample). For all panels except I: *, P ≤ 0.05, **, P ≤ 0.01, ****, P ≤ 0.0001 as assessed by Student’s *t* test. Panel I: ****p <0.0001 assessed by two-way analysis of variance (ANOVA). Arrows indicate micronuclei (MN), nucleoplasmatic bridges (NPB) and nucleoplasmatic bud (NBUD).

These data supported the hypothesis that USP15 depletion would render normal hematopoietic progenitors more sensitive to genotoxic stress *in vivo*. To test if this is the case, we injected mice with the chemotherapeutic agent cisplatin (0,8 mg/kg, a relative low dose compared to the 6mg/kg maximal tolerable dose used in C57BL6/6J mice (Pilzecker et al., 2017)) i.v or with PBS, and analyzed the BM after 2 days. Upon cisplatin treatment, USP15 knockout BM cells produced significantly fewer CFUs compared to wt (Fig. S6I), suggesting higher sensitivity of their HSPC compartment. Deeper BM analysis unmasked a broader sensitivity of the primitive progenitor compartment in *Usp15-/-* mice, including HSCs and LSKs, and also the more proliferative LKS-, myeloid (GMP) and lymphoid (CLP) progenitor populations to genotoxic stress (Fig. S6J, K). This effect may also be seen as an orthogonal validation of our results obtained upon transplantation-induced (replication) stress, overall suggesting that USP15 generally contributes to preserve the HSPCs compartment during stress-induced hematopoiesis.

Finally, we sought to translate these findings into a potential combination setting in leukemia. In leukemia cells originated by blast crisis such as KBM7 cells, we combined depletion of USP15 by dox-inducible RNAi and DNA single/double-strand breaks induction by ionizing radiation (IR). USP15 depletion by a dox-inducible shRNA sensitized KBM7 cells to IR (Fig. 7H). A role for USP15 in supporting the DNA damage response was recently reported in breast cancer cells (Peng et al., 2019). In keeping with those findings, Rad51 protein levels were diminished by USP15 knockdown in MV411 and Kasumi-1 leukemia cells (Fig. S6L). A broader chemo-profiling in CCLE cancer cell lines indicated that leukemia cell lines are generally more sensitive than others to the DNA damage inducers Topotecan (Topoisomerase I inhibitor) and Mitomycin-C (a DNA cross-linking agent), two chemotherapeutic clastogenic agents (Fig. 7I). Notably USP15-depletion cooperated with Mitomycin-C to reduce cell viability in MV4-11 (Fig. 7J).

Overall, these data suggest that USP15 generally safeguards genome integrity in normal hematopoietic and leukemia cells and its loss affects survival in a selected subset of leukemia cells and sensitizes these cells to DNA damage inducers.

## Discussion

We report on the comprehensive assessment of the role for deubiquitinating enzymes in early hematopoiesis through the successful application of pooled *in vivo* shRNA screens in the mouse. Through focused *in vivo* RNAi screens, we have identified several genes within the family of DUBs whose loss increases or decreases mouse hematopoietic stem and progenitor cells fitness *in vivo.* Remarkably, in this unbiased approach, several known and novel DUBs were shown to regulate hematopoietic homeostasis. We followed upon USP15, which we characterized as a novel DUB required for early hematopoietic progenitor proliferation and for HSC homeostasis *in vivo*. Also, our findings uncover a conserved requirement for USP15 in preserving normal and leukemic stem cell genome integrity.

Pooled *in vivo* screens in early progenitors pose specific technical challenges related to the heterogeneity of the starting population, the rare set of target stem and progenitor populations, the (long-term) grafting potential, and to selective pressure *in vivo*. The success of our shRNA screening approach is underscored by the successful shRNA library representation in Lin-cells, engraftment of transduced cells, detection of a large library representation *in vivo* in Lin- and in splenic B cells (≥90%) and retrieval of positive shRNAs controls *in vivo*. Also, shRNAs targeting DUBs previously involved in HSC biology were scored in the screens, in particular USP1 (Parmar et al., 2010), USP3 (Lancini et al., 2014), USP16 (Adorno et al., 2013; Gu et al., 2016) and BAP1 (Dey et al., 2012; Higuchi et al., 2016; Nakagawa et al., 2015; Nijnik et al., 2012; Wang et al., 2013) (Table S3). Together with the extensive genetic validation, these examples raise confidence in the identification of USP15 as critical regulator of HSCs *in vivo* and validate the use of *in vivo* genetic screens for systematically characterizing the role of DUBs in tissue homeostasis.

Loss of USP15 in adult murine hematopoietic progenitors by RNAi or by germline deletion impaired their growth *in vitro* and repopulation ability *in vivo*. In USP15 knockout adult mice, HSC reserves in particular were significantly reduced, while more differentiated progenitors were largely maintained. These data suggest that chronic lack of USP15 likely allows for protective pathways/adaptation that ensure hematopoiesis at steady state and are consistent with a role for USP15 in contributing to homeostasis through the maintenance of HSCs, which are largely quiescent (Bakker and Passegue, 2013). HSC regenerative potential is, in fact, critical when HSC are induced into proliferation during *ex-vivo* culturing, (competitive) transplantation (Flach et al., 2014; Walter et al., 2015), or upon aging (Beerman et al., 2010; de Haan and Lazare, 2018).

We report that spontaneous genotoxic stress and enhanced sensitivity to clastogenic agents accompanied the decrease in viability of USP15 deficient hematopoietic progenitors and leukemia cells *in vitro* and of mouse primitive hematopoietic progenitors *in vivo*. These data link USP15 to the DDR and are consistent with previous work in cancer cell lines (Fielding et al., 2018; Mu et al., 2007; Nishi et al., 2014; Peng et al., 2019). Although a genome maintenance defect may contribute to loss of fitness of USP15 deficient HSCs, understanding how USP15 modulates the damage response and how its loss precisely impacts on HSC and cancer cells maintenance will require deep molecular characterization, not least through the identification of context-specific interactors. This endeavor may be facilitated by the current development of single-cell proteomics approaches.

USP15 gene alterations have been found in cancer, in particular glioblastoma, breast, ovarian and recently in non-small cell lung cancer (Eichhorn et al., 2012; Fielding et al., 2018; Padmanabhan et al., 2018; Peng et al., 2019), as well as copy number losses in pancreatic cancer (Srihari and Ragan, 2013). Here, the analysis of cancer databases uncovers USP15 highest expression in primary blood-derived tumors and alterations in leukemia patients, particularly CML and AML. This is the first study to provide functional ground for investigating the role for USP15 as gatekeeper in leukemia. Our *in vitro* assays support targeting USP15 to suppress viability of a subset of leukemia cell lines, including cell lines derived from blast crisis stage CML, a stem-cell derived disease (Arrigoni et al., 2018). In both normal and tumor cells, USP15 inhibition is accompanied by chromosomal aberrations indicating a susceptibility to DNA damage. Further research is needed to examine the role of USP15 in human HSC and the impact of USP15 loss of function or inhibition on steady-state human hematopoiesis.

USP15 in involved in multiple cellular processes, including p53 (Liu et al., 2017; Zou et al., 2014) and NF-kB (Schweitzer et al., 2007) signaling, histone H2B deubiquitination (Long et al., 2014), and the transforming growth factor-β (TGF-β) pathway (Eichhorn et al., 2012; Inui et al., 2011; Iyengar et al., 2015), and showed interaction with a number of ubiquitin linkages *in vitro* (Zhang et al., 2017). Interestingly, recent studies have linked USP15 to inflammation in experimental models, through the regulation of type I and type II interferon (IFN) responses (Pauli et al., 2014; Torre et al., 2017; Zou et al., 2014; Zou et al., 2015). Also, by diminishing TGF-β signaling, depletion of USP15 was shown to reduce the proliferation and the oncogenic capacity of patient-derived glioma-initiating cells (Eichhorn et al., 2012). TGF-β regulatory mechanisms and inflammatory signals are relevant both in normal HSC and in malignant development (Blank and Karlsson, 2015). Notably, upon USP15 knockdown in CML lines, we find significant changes in gene expression including up-regulation of genes associated with inflammation and down-regulation of TGF-beta signaling genes, suggesting that USP15 role in these cells may involve alterations in these pathways. While our data in leukemia cell lines indicate that context-specific pathways are likely to be affected by loss of USP15 in cancer cells, notably TGF-beta and inflammatory pathways, the limited changes in gene expression detected in LSK cells of the USP15 knockout mice suggests that Usp15 function in preserving genome integrity is dominant in this compartment. This may open to potential cancer cell specific drug combinations, which would spare normal cells from side effects.

The function of USP15 in development is still poorly characterized. Here, we demonstrate a requirement for USP15 in HSC maintenance. Our results also support a broader role of USP15 in mammalian development, as knockout of USP15 has clear impact on Mendelian transmission and on mouse lifespan. This finding is not obvious when compared against previous reports in a USP15 gene-trap model (Zou et al., 2014). In our study, germline deletion of the entire USP15 locus by CRISPR/Cas9 generates mice with a complete gene knockout, allowing the manifestation of the full phenotype at organismal level. Growth retardation observed in our USP15 knockout mice is, in fact, in line with recently reported findings in another CRISPR-Cas9 USP15 knockout mouse model (Peng et al., 2019). Our mouse model enriches the toolbox available to the quest for mechanisms underlying USP15 function *in vivo*.

USP4, USP11 and USP15 are part of a closely related family of USPs (Al-Salihi et al., 2012; Das et al., 2017; Eichhorn et al., 2012; Long et al., 2014; Nishi et al., 2014; Orthwein et al., 2015; Vlasschaert et al., 2015; Wijnhoven et al., 2015; Zhang et al., 2012; Zhou et al., 2017). They are all expressed in hematopoietic early progenitors (Cabezas-Wallscheid et al., 2014; Heng et al., 2008; Lancini et al., 2016), but none was previously linked to HSC activity. All three genes scored as hits in our screen, suggesting that besides USP15, its paralogues may contribute to HSC homeostasis and might functionally cooperate. Notably, USP15 deficiency in mice was suggested to be lethal when the mice were bred with USP4-targeted mice, suggesting partially redundant roles *in vivo* (Vlasschaert et al., 2015). The discovery of USP15, USP11 and USP4 individually as critical modulators of HSC homeostasis opens to the study of specific and redundant roles in a physiological setting. The potential redundancy and biochemical interaction between USP15 and USP11 may also open to the rational design of allosteric degraders, which would have a stronger impact than individually targeted small molecules.

In summary, we employed an unbiased approach to sensitively and selectively screen for DUB function in hematopoietic progenitors *in vivo,* through which we identified known and novel DUB candidates. Major investments are being devoted in the last 5-10 years to DUB drug discovery, with more than 40 small molecules against DUBs already developed (Harrigan et al., 2018; Heideker and Wertz, 2015). Our data strongly argue in favor of developing specific USP15 inhibitors, which are only starting to emerge. Interestingly, engineered ubiquitin variants targeting USP15 have recently been reported (Teyra et al., 2019), which may open up to the development of more specific small molecules as well as targeted degraders. More broadly, our study calls for a more systematic effort in understanding how DUBs regulate normal and malignant HSC biology as a critical route towards the selection of effective drug targets and targeted treatment combinations.

## Acknowledgements

We thank T. Sixma for critical reading of the manuscript, R. Kerkhoven, I. de Rink and W. Brugman from the NKI Genomic Core Facility for deep-sequencing, A. Pfauth, F. van Diepen, and M. van Baalen for assistance during flow cytometry, G. Chandrasekaran for data visualization, F. Alemdehy, the MCCA transgenic facility, B. van den Broek and the NKI BioImaging Facility, the NKI Intervention Unit, the animal facility and Animal Pathology Department for assistance. Drug sensitivity data were taken from the CTD2 Network (https://ocg.cancer.gov/programs/ctd2/data-portal) established by the National Cancer Institute’s Office of Cancer Genomics. This study was supported by the Dutch Cancer Society (KWF) grant to E. Citterio and M. van Lohuizen (NKI-2012-5665 grant) and by the ONCODE Institute to M.v.L.

## Author contributions

Conceptualization and validation: E.C, G.G.; investigation and methodology: E.C., P.v.d.B., C.L., C.C., D.H., M.S., JY.S., M.J.S., G.G.; resources: P.v.d.B., C.C., M.S., C.P., JY.S.; I.J.H., H.J.; formal analysis: P.v.d.B., C.L., C.C, G.G., E.C.; software: C.C., G.G.; writing-original draft: E.C.; writing-review and editing G.G., P.v.d.B, H.J., M.S.; supervision and funding acquisition: E.C., G.G., M.v.L. All authors read and commented on the manuscript.

## Materials and Methods

### EXPERIMENTAL MODELS

#### Mice: generation and breeding of USP15 knockout mice

Full *Usp15* knockout (KO) mice (MGI: Usp15<em1Nki>; MGI:5810631; B6J-Usp15<em1Nki>) were generated by CRISPR-Cas9-mediated deletion of the *Usp15* locus in C57BL/6J zygotes as described (Pritchard et al., 2017). Two CRISPR-guides (gRNAs) were used that target Cas9 mediated double stranded DNA cleavage at both the 5’ and 3’ UTR of Usp15. The sequence of the 5’ and 3’ targets were TCTTCTTCCACTAGCC<u>GT</u>AGCGG and GTCACTTGATACGATA<u>GC</u>GCCGG, respectively. The expected cleavage sites (underlined) are 91.795 bp apart in the C57BL/6J genome (Fig. S4A). Mice carrying a full *Usp15* knockout (KO) allele, in which the 91 kb of cleavage site intermitting sequence is missing, were identified by PCR and sequence analysis. The *Usp15* alleles were detected with the following primers: forward, 5’-TCCAGTAGGAGTGAACCCGC-3’; reverse KO, 5’-AGGTGGCTGAGAGTGAGAGCAGG-3’; reverse wt, 5’-GCCTTCCGCCATCTTCTTCCAC-3’, yielding a product of 590 bp and 396 bp for *Usp15*-KO or *Usp15* wt, respectively. The predicted CRISPR-Cas9-mediated fusion product for the *Usp15*-KO allele is: ccgcta.c.tatcgtat. The 590bp *Usp15-KO* PCR fragment was sequenced, yielding the obtained fusion product: ccgtaTcGGatcgtat. Genotyping was performed by PCR of genomic tail DNA using the Extract PCR kit (Bioline, cat. No. BIO-21127). All mice were kept on C57BL/6J (The Jackson Laboratory) strain background (CD45.2^+^) in a specific pathogen-free environment. *Usp15* mice were maintained heterozygous. All animal experiments comply with Dutch and European regulations and ethical guidelines and have been authorized by our local experimental animal committee at The Netherlands Cancer Institute (DEC-NKI). For determining survival, mice time to death was defined as the latency between birth and unexpected death or a terminal disease stage indicated by >20% weight loss or other symptoms of severe sickness. Mice were sacrificed by CO_2_ asphyxiation and underwent necroscopy. Organs were collected and fixed for histopathological analysis as described (Lancini et al., 2014).

#### Cell lines and cell culture

##### Leukemia cells

KBM7 were grown in IMDM (Gibco) medium. All the other leukemia cell lines, including K562 and MV4-11, were grown in RPMI 1640 (Gibco) medium. Culture medium was supplemented with 2 mM L-Glutamine (Gibco 25030-164), 10% fetal calf serum (FCS, Thermo Scientific) 100 U ml^−1^ penicillin, and 100 µg ml^−1^ streptomycin (Pen/Strep Gibco 15140-163). Cells were incubated at 37°C in a humidified atmosphere containing 5% CO_2_.

##### U2OS cell line

U2OS osteosarcoma cells were grown in Dulbecco’s modified Eagle medium (DMEM; Gibco), supplemented with 10% fetal calf serum (FCS, Thermo Scientific), 100 U ml^−1^ penicillin, and 100 µg ml^−1^ streptomycin (Pen/Strep Gibco 15140-163). Cells were incubated at 37°C in a humidified atmosphere containing 5% CO_2._

### METHOD DETAILS

#### shRNA libraries

Lentiviral hairpins (pLKO.1) targeting annotated DUB genes(Mevissen and Komander, 2017) and controls were selected (Table S1). Vectors were individually picked from glycerol stocks of The RNAi Consortium library (TRC Mm1.0) (Sigma-Aldrich, MO), grown up on agar plates and combined before maxiprep DNA isolation. Pooled plasmid libraries were used to produce lentiviral particles using standard procedures (Gargiulo et al., 2014).

#### *In vivo* shRNA screens

Lineage negative (Lin-) hematopoietic stem and progenitors cells were isolated from the bone marrow (BM) of 8-10 weeks old wild-type (wt) mice (C57BL/6J-Ly5.2)(CD45.2) and plated in serum-free medium supplemented with cytokines as described below. Lin-cells were pre-stimulated for 24 hours (hrs) and transduced with pooled lentiviral shRNAs at low multiplicity of infection (MOI <0.5) using spin-inoculation at 1,800 rpm for 90 min at 32 °C. Twenty-four hours after infection, the cells stably expressing integrated shRNA were selected with puromycin (1μg/ml). 48 hours later, cells were harvested, keeping 1 x 10^6^ cells for DNA extraction of the INPUT(T0) sample. Next, 2×10^6^ Lin-cells for the primary screening or 1×10^6^ for the secondary screens (representing a minimum of 3,500-fold enrichment over the library), were mixed with 1×10^6^ freshly isolated BM cells from wt (C57Bl/6-Ly5.1)(CD45.1) mice and injected into recipient mice as described below. Recipient mice were sacrificed at 4 weeks post transplantation (wpt) and femurs, tibia, and spleen were collected. Lin-cells from BM and CD43-, CD19+, CD220+, CD45.2 splenocytes were purified as described below and genomic DNA was extracted for PCR amplification of the shRNAs.

#### PCR and next-generation sequencing

shRNAs sequences were retrieved from genomic DNA by PCR amplification as described (Gargiulo et al., 2014). For every sample, a maximum of 8 µg genomic DNA was divided over 4 50-µl PCR reactions using barcoded forward primers (PCR1). The products of all reactions were pooled and a maximum of 1 μg from this PCR1 was used per reaction in subsequent PCR2 reactions using primers containing Indexes for next generation sequencing. Barcodes and Indexes for deep sequencing (Illumina HiSeq 2000) were incorporated into PCR primers as listed in Table S4.

PCR mixture per reaction: 10 µl HF Buffer (NEB), 1.25 µl 10-µM forward primer, 1.25 µl 10-µM reverse primer, 1.25 µl Phusion Hot Start II polymerase (2U μl^-1^; Thermo Scientific, cat.n. F-530L), 1 µl 10-mM dNTPs, DMSO 3% (vol/vol), adding mQ and template to 50 µl. PCR conditions were: 1’ @ 98 °C, 16 (PCR1) or 14 (PCR2) x (10s @ 98 °C, 30s @60 °C, 60s @ 72°C), 5 min @ 72 °C. PCR products were purified using the AMPure XP solid-phase reversible immobilization kit (SPRI; Beckman Coulter, cat. no. A63881) and subjected to Illumina next-generation sequencing. The shRNA sequence reads were aligned to the TRC library. Fold change in individual hairpin representation *in vivo* was determined by comparing shRNA representation in each sample to that in the control cell population remaining after tail vein injections during bone marrow transplantation (INPUT, T0). Each condition included in the preliminary analysis was matched to its corresponding shRNA library removing those shRNA that weren’t present in any of the samples. Pairwise differential abundance analysis was performed between test sample and input using limma v3.36 (Ritchie et al., 2015) after outlier removal using PCA. shRNA were considered as enriched or dropped out if logFC was higher than |1|, adj.Pvalue <= 0.02 and avg. abundance > 2.5 . The analysis was done using R v3.5 programming language (https://cran.r-project.org/).

#### Bone marrow transplantation assays

For RNAi *in vivo* screens, puromycin selected, retroviral-transduced Lin-cells (CD45.2) were mixed with wt bone marrow cells (CD45.1) as described above and injected into lethally irradiated (2 doses of 5.5 Gy TBI separated by an interval of 3 hours) wt C57BL/6-CD45.1 recipient mice. Donor contribution was assessed based on the expression of CD45.1/CD45.2 antigens. At 4 wpt primary recipients were sacrificed and the frequency of donor-derived CD45.2 peripheral blood cells, splenocytes, Lin- and LSK were assayed by phenotypic profiling. Lin-cells and CD45.2 splenic B cells for genomic DNA extraction and shRNA retrieval were isolated as described below.

In validation experiments, wt Lin-cells were transduced with individual lentiviral vectors and puromycin-selected as indicated above. 1x10^6^ lentiviral-transduced, puromycin resistant cells (CD45.2) were transplanted together with 1x10^6^ total BM cells (CD45.1) into lethally irradiated recipient mice (CD45.1). Recipient mice peripheral blood was monitored by FACS analysis at 2 wpt and every 4 wk for 18 wk. Donor contribution and multilineage reconstitution were assessed based on the expression, respectively, of CD45.1/CD45.2 antigens or CD19, CD3, and Gr1 markers in the CD45.2^+^ fraction. At 18 wpt, primary recipients were sacrificed and the frequency of donor-derived CD45.2 B and T cells in the spleen and Lin-, LSK, HSC and myeloid progenitors in the bone marrow was assayed by phenotypic profiling.

In competitive BM transplantation, BM was isolated from donor test animals (CD45.2, *Usp15+/+* or *Usp15-/-*) and mixed in a 1:1 ratio with wt competitor cells (CD45.1)(8-10 weeks old mice). For each genotype and for the wt competitor BM, cells from 3 donors and from 3 wt competitor mice were isolated and pooled before 1:1 mixing. Thereafter, 1 × 10^6^ CD45.1/CD5.2 mixed BM cells were transplanted into lethally irradiated CD45.1 recipient mice as described above. Flow cytometry staining for LSK and HSC of donors was performed to ensure that the HSC frequency in test and control BM would be comparable. Differences in HSC were corrected before transplantation to transplant stem cell equivalents. Chimerism in the blood of primary recipients and BM repopulation at 18 wpt were assessed as described above.

Mice were irradiated using Faxitron MultiRad 225 X-ray irradiation system. Irradiated mice were treated with Enrobactin for the first 4 wk after irradiation. Immunophenotyping, Lin-isolation and CD45.2 splenocytes purification were performed as described below.

#### Flow cytometry

##### Blood

Peripheral blood was collected into EDTA-coated micro-tubes. For FACS analysis, blood was depleted from red blood cells by hypotonic lysis and staining was performed with fluorochrome-labelled antibodies, CD3-FITC, CD11b-PerCp/Cy5.5, CD19-APC, Gr1-APC/Cy7.

##### Spleen

Single cell suspension of spleen was obtained by smashing through a 70μm filter. Suspension was depleted from red blood cells by hypotonic lysis. For isolation of immature and of mature (resting) B cells, CD43 positive cells were first depleted using anti-mouse CD43 (Ly-48) MicroBeads (MACS Miltenyi Biotec). Donor derived cells were then isolated by FACS sorting using fluorochrome-labelled antibodies CD45.2-FITC, CD45.1-PE, CD19-APC, B220-Pb, CD43-biotin and Streptavidin APC/Cy7. For immunophenotyping of spleen upon transplantation, staining was performed with fluorochrome-labelled antibodies, CD3-FITC, CD8a-PerCp/Cy5.5, CD4-APC, CD19-APCH7.

##### Analyses and cell sorting of hematopoietic precursors

To analyze Lin-, LSK, HSC, LKS-, CMP, GMP and MPP subpopulations (Cabezas-Wallscheid et al., 2014; Kiel et al., 2005; Oguro et al., 2013; Wilson et al., 2008; Yeung and Wai Eric So, 2009), BM freshly isolated mononuclear cells (MNC) were first stained with Lineage Cell Detection Cocktail-Biotinylated mouse antibody (MACS Miltenyi Biotec). For FACS analysis, cells were then directly stained with fluorochrome-conjugated antibodies. We used 5 x 10^6^ MNCs per staining. For quantifying LSK and HSC populations, cKit-APC, Sca-1-PerCp/Cy5.5, CD48-FITC, CD150-PE/Cy7, CD135-PE and streptavidin-APC/Cy7 antibodies were used. For quantifying LKS^-^ progenitor populations, cKit-APC, Sca-1-PerCp/Cy5.5, CD34-FITC, CD16/32-PECy7, and streptavidin-APC/Cy7 antibodies were used. For purifying Lin-cells for shRNA library viral infection or for culturing, depletion of lineage^+^ cells from MNCs was performed using Biotin labeled Lin+ cocktail and Streptavidin MicroBeads (Macs; Miltenyi Bio-tec) and magnetic columns (Macs; Miltenyi Biotec). For cell sorting of LSK, depletion of lineage^+^ cells was first performed as above before staining.

##### Cell cycle analysis of BM populations

Cell surface staining was performed as described above. Samples were then fixated in 3.7% Formaldehyde in PBS (Sigma)) for 30 min at RT. Cells were permeabilised in PBS/BSA(1%)/Tween20 (0.025%)(PBT) for 15min at RT and harvested in PBT containing 10 **µ**g/mL DAPI for chromatin labeling. Cell cycle analysis was performed as described (Pilzecker et al., 2017).

##### Assessing Cleaved Caspase-3 levels of BM populations

Cell surface staining, fixation and permeabilization was performed as described above followed by staining with Cleaved Caspase-3-AF488 antibody for 30 min at RT in PBT. Cells were washed twice with PBT and harvested in PBT containing 10 **µ**g/mL DAPI for chromatin labeling.

All FACS measurements were performed with a BD LSRFortessa cell analyzer (BD Biosciences). Cell sorting was performed with a FACSAria (BD). All FACS data were analyzed using FlowJo Software version 10.0.8r1. (Tree Star).

#### Hematopoietic stem and progenitor cells liquid culture, time-lapse imaging and proliferation assays

Mice were sacrificed at the indicated age (8-12 weeks). Lineage negative (Lin-) isolation, LSK FACS sorting and cell surface staining was performed as described above. Cells were plated on Ultra-Low Attachment multiwell plates (Corning→Costar→) in StemPan SFEM (*StemSpan*™ Serum-Free Expansion Medium (*SFEM*) STEMCELL Technologies) supplemented with mouse SCF 100 ng/ml, mouse thrombopoietin (mTpo) 50 ng/ml and mFlt3 ligand 50 ng/ml (PrepoTech)(Ye et al., 2008). Medium was replenished and cells were expanded in 3% oxygen to maintain optimal growth. To evaluate proliferation, 1,000 Lin-cells were plated in 96-well plates at day 7 of culture. 4 wells per conditions were imaged (phase-contrast) with a 4 hrs interval for 6.5 d using the IncuCyte FRL (Essen BioScience). Confluence was determined by the IncuCyte software, based on area (confluence) metrics. Plating of 500 cells/well gave similar results. FACS sorted LSK cells from individual animals were grown individually. LSK were plated at day 8 of culture and monitored for growth by counting live cells by Trypan blue exclusion using a TC20^TM^ Automated Cell Couter (BIORAD) at the indicated time. Four wells per condition were counted.

#### CFU-C colony-forming assay

BM MNCs cells were seeded on 35-mm culture dishes in triplicate in methylcellulose medium supplemented with cytokines (MethoCult GF M3434, STEMCELL Technologies). CFU-Cs (colony forming units in culture) include CFU-GEMM (granulocyte, erythroid, macrophage, megakaryocyte), multipotential progenitors and lineage-restricted progenitors of the erythroid (BFU-E, burst-forming unit–erythroid), and granulocytic, monocyte-macrophage (CFU-GM). Cultures were incubated at 37°C under 5% CO_2_. Colonies were quantified at day 8.

#### Leukemia cells RNA interference and cell viability assays

USP15 siRNAs (siGENOME Human USP15 (9958) siRNA-SMART pool M-006066-01) and control (Ctrl) siRNAs (siGENOME Non-Targeting siRNA Control Pool#2 D-001206-14-05) were from Dharmacon. Cells were transfected using Lipofectamine→ RNAiMAX Reagent from Life Technologies following the manufacturer’s instructions and assayed at 72 hrs after transfection in Western blotting, viability assays and immunofluorescence or RNA-seq.

For inducible USP15 knockdown experiments, we have tested 4 shRNAs. To rule out potential off-target effects by one shRNA, we used two different shRNAs for experiments in Figure 7H-J (sequences ‘TAAACCAGCATCCTGAATGG’ and ‘TTTCATGAACTCAGCTATTC’, respectively). The additional shRNA tested were: ‘GCATTAGGCTGCCGTATATA’, and ‘CGCTTATAAGAACTATGATT’ and these were found insufficiently potent on-target. The shRNA-containing bicistronic vectors were either subcloned in synthetized at VectorBuilder, and included a H1 promoter and Tet operator shRNA cassette as well as a Tet repressor-T2A-eGFP for FACS sorting of TetR positive cells.

Viability assays were performed using AlamarBlueTM Cell Viability reagent (Thermo Fisher Scientific). Relative viability was normalized to the control siRNA transfected cells and corrected for back-ground signal.

#### TCGA pan-cancer gene expression analysis

Gene expression analysis by RNAseq was compiled using data from all TCGA cohorts. Gene expression was measured using the IlluminaHiSeq technology. Data from all TCGA cohorts are combined to produce this dataset. Values are PANCAN expression unit - (log(norm(exp) + 1)) transformed RSEM values.

#### RNA-seq gene expression analysis

For gene expression analysis, KBM7 or K562 cells were transfected with USP15 or control siRNAs as described above. Total RNA was extracted at 72 h after transfection. For LSKs, cells were FACS sorted from freshly isolated BM and total RNA was extracted. n=3 *Usp15+/+* and *n=2 Usp15-/-* littermates (2 months old). Samples were prepared using TruSeq protocols, and standard sample preparation protocols and RNA-seq was performed on a Hiseq2000 machine (Illumina) at the NKI Genomics Core Facility. The data discussed in this publication will be available from the NCBI’s Gene Expression Omnibus database (Edgar et al., 2002).

#### Sensitivity assays

Cells were cultured in the proper culture medium with doxycycline for 5 days (100ng/ml) to induce USP15 knockdown. Cells were then seeded in 384 well plates 24 hrs before treatment. For IR, cells were irradiated with the indicated dose and cell viability was assessed 3 days after IR using a medical irradiator platform (XenX). To determine the 30nM MMC concentration, MMC dose-response experiments were previously assessed and 30nM was determined as the IC50 dose for MV4-11 cells in our experimental conditions. Cell viability was assessed at 72 hours. Doxycycline and drug were refreshed daily. In vivo cisplatin (CsPt) Sensitivity Assay. Mice were injected i.v. with 0.8 mg/kg cisplatin or PBS. After 2 d, the BM was isolated and analyzed as described above.

#### Immunofluorescence and quantitative image analysis

*In vitro* cultured murine LSK or human K562 and KBM7 cells were deposited on charged slides (Superfrost Plus; Menzel-Glaser) by cytospin and directly fixed in 4% paraformaldehyde. Immunostaining with antibodies against 53BP1 was performed as previously described (Lancini et al., 2014). Counterstain was with Alexa Fluor 488–conjugated secondary antibodies and DAPI (200 ng/ml) (Sigma-Aldrich). Micronuclei were scored on fixed cells stained with DAPI. Digital images were acquired using a microscope (AxioObserver Z1; Carl Zeiss) with an ORCA-ER CCD charge-coupled device) camera (C4742-80-12AG; Hamamatsu) and Zen software (40x and 63x magnification). A macro in ImageJ software (version:2.0) (developed by Bram van den Broek, NKI, Amsterdam) was used for quantification of spontaneous 53BP1 DNA damage foci. The DAPI channel was used to select the nuclei of the cells in the field. Briefly, Z-stacks are converted to two dimensional via one of several user-defined methods: maximum intensity projection, automatically select sharpest slice or manually select a slice. Region of interests (ROIs) of candidate nuclei are then automatically obtained throughout the image stack by auto-thresholding an outlier-removed median-filtered (0.7 mm radius) *z* projection of the nuclei channel, followed by a watershed command to separate touching nuclei and particle analyser run with size (>4 and 40 μm2), and circularity (>0.25) constraints. In the detection of 53BP1 foci, the foci threshold level is defined by the signal-to-noise ratio (SNR): a (user-set) factor times the s.d. of the background fluorescence intensity of the nucleus. The latter property is approximated by first crudely removing signal outliers (the foci), and then taking the median and s.d. of the lower approximate 80% pixel values in the ROI, respectively. The background intensity is subtracted using a Difference of Gaussians filter. Foci are then identified as regions of adjacent pixels with grey values, exceeding the SNR threshold and area larger than a certain minimum. In the procedure, the SNR is the only user-defined parameter, and is iteratively optimized by comparing the detected foci with the original signal in an overlay image.

#### Protein analysis

Cell were lysed with RIPA buffer (10 mM Tris-Cl (pH 8.0), 1 mM EDTA, 1% Triton X-100, 0.1% sodium deoxycholate, 0.1% SDS, 140 mM NaCl, 1 mM PMSF) containing protease inhibitor cocktail (Complete, Roche) and phosphatase inhibitors 9Na Fluoreide 10mM final concentration, Na orthovanadate 1 mM final concentration and NaPPi 1 mM final concentration) and whole cell extract was loaded on SDS-PAGE on NuPAGE gels, followed by Western blotting with the indicated antibodies (Table S4). Filter blocking and antibody incubation were performed in PBS supplemented with 0.1(v/v) % Tween and 5%(w/v) bovine milk powder.

#### Quantitative real-time (qRT) PCR

Total RNA was extracted using TRIZOL reagent (Life Technologies) or ReliaPrepTM RNA miniprep System (Promega) and cDNA was prepared using Superscript II RT and oligo(dT)_n_ primers (Life technologies). qRT-PCR was performed on a StepOnePlusRT-PCRsystem(AppliedBiosystems) usingSYBR Green PCR Master Mix (Applied Biosystems). The amount of target, normalized to an endogenous reference (HPRT), was calculated by 2^-ΔΔC^T. Primer sequences were as follows: *Hprt* forward, 5’-CTGGTGAAAGGACCTCTCG-3’; *Hprt* reverse, 5’-TGAAGTACTCATTATAGTCAAGGGCA-3’; *Usp15* forward A, 5’-TGTGGCTTAAGTAACTTGGGAAA-3’; *Usp15* reverse A 5’-AAGTGGAGGTGTGTTGCTCA-3’; *Usp15* forward B, 5’-TCAGCTGGTACACACTGATGG-3; *Usp15* reverse B, 5’-TGCTTTACAAACATACCCTGTTCT-3’. Primers used in Fig. S5G,H for validation of RNA-seq results are available upon request.

#### Antibodies

Antibody specifications are listed in Table S5 and S6.

#### sgRNA design and cloning

sgRNA sequences targeting human USP15 were selected from the Human Brunello CRISPR knockout pooled library (Doench et al., 2016)(Addgene #73178) and further selected on the basis of high quality score in two additional online tools: CRISPR Design (http://crispr.mit.edu/) and CRISPRscan (http://www.crisprscan.org/). The following sgRNA (sg01) targeting exon 3 of USP15 DUSP domain was used in this study: *USP15ex*3, 5’-AAGGTGTTCCTTAAGTGACT-3’. Pairs of complementary DNA oligos (for: 5’-caccgAAGGTGTTCCTTAAGTGACT-3’; rev: 5’-aaacAGTCACTTAAGGAACACCTTc-3’) were annealed and the DNA oligonucleotide duplex was cloned in the Bbs1 restriction site of pSpCas9(BB)-2A-GFP (PX458) plasmid (Plasmid #48138, Addgene). sgRNA sequence was verified by DNA Sanger sequencing using the following primer: *ASK*-*FN2* 5’-CGGCCTTTTTACGGTTCCTG-3’.

#### Transfection and fluorescence activated cell sorting (FACS)

Plasmids were transfected into U2OS cells using FuGENE® HD Transfection Reagent (Promega), according to the manufacture’s instructions. After 48 hours in culture, cells were trypsinized, washed with PBS, and resuspended in PBS (supplemented with 3% BSA) and passed through a cell strainer (*Falcon*® Round-Bottom Tubes with *Cell Strainer Cap,* Catalog #38030). Cells were individually sorted (BD FACSAria, BD FACS Diva 8.0.1 sofware) based on EGFP signal into tissue culture 96-well plates (CELLSTAR-Greiner) at a single cell per well for clonal expansion. Viable individual clones were then transferred to 24 well plates for clonal expansion and screening.

#### Genomic DNA extraction, PCR and Sanger sequencing and TIDE analysis

Genomic DNA was extracted using DNeasy Blood & Tissue Kit (QIAGEN). The genomic region around the gRNA-targeted sequence was amplified using the following primers: for,5’-GTTAGTGTTACAATTCTTCCAATACGG-3’; rev,5’GTTTTATCAAAAACAGTGCAGCACAG-3’. PCR was performed using Thermo Scientific^®^ *Phusion* High-Fidelity DNA polymerase in GC buffer. PCR conditions were as follows: 30s at 95°C, followed by 325 cycles of 30s at 95 °C, 30s at 64 °C, and 30s at 72 °C, followed by 3 min at 72 °C. Primer 5’-TTACAATTCTTCCAATACGGCCCAG-3’ was used for Sanger sequencing. About 100-200 ng DNA from purified PCR samples was prepared for sequencing using BigDye terminator v3.1. Samples were analyzed by an Applied Bio-systems 3730×1 DNA Analyzer. The data obtained was analyzed using the TIDE software (https://tide.nki.nl). The decomposition window used for TIDE was set to indels of size 0-10 bp, p threshold of 0.001.

### QUANTIFICATION AND STATISTISAL ANALYSIS

Statistical analysis was performed in GraphPad Prism 7.0, using an unpaired two-tailed Student’s *t* test or Multiple t test. Animal survival experiments were analyzed with a Log-rank nonparametric test and expressed as Kaplan-Meier survival curves. In all Fig.s: *, *P* <0.05; **, *P* < 0.01; ***, *P* < 0.001; ****, *P* < 0.0001.

### DATA AND CODE AVAILABILITY

The RNA-sequencing data discussed in this publication will be available from the NCBI’s Gene Expression Omnibus database (Edgar et al., 2002).

## Abbreviations

AML, acute myeloid leukemia; CML, chronic myeloid leukemia; CMP, common myeloid progenitor; DDR, DNA damage response; DSB, DNA double-strand break; DUB, deubiquitinating enzyme; GMP, granulocyte-monocyte progenitor; HSC hematopoietic stem cell; Lin-, lineage negative; LSK, Lin^−^Sca1^+^c-Kit^+^; LKS^-^, Lin^−^Sca1^-^c-Kit^+^; LSC, leukemic stem cell; MEP, megakaryocyte-erythrocyte progenitor; MNC, mononuclear cell; USP15, ubiquitin-specific protease 15; PBC, peripheral blood cell; wpt, weeks post transplantation.

## Supplemental Information

### Supplemental Figures

Figure S1 (relative to Figure 1). *In vivo* RNAi screens for DUBs in murine hematopoietic stem and progenitor cells.

Figure S2 (relative to Figure 1). Pooled *in vivo* RNAi screens identify candidates DUBs effectors of HSC activity.

Figure S3 (relative to Figures 2C, 3, 4 and 5). Flow cytometry and gating strategies.

Figure S4. Generation and initial characterization of *Usp15* knockout mouse model (relative to Figure 4).

Figure S5. USP15 RNAi in CML cell lines results in alteration of inflammatory signaling pathways.

Figure S6. USP15 loss enhances genotoxic stress *in vitro* in human cancer cells and *in vivo* in mouse hematopoietic stem and progenitor cells.

### Supplemental Tables

Table S1 (relative to Figure 1). Genes targeted and shRNAs used in the primary and secondary *in vivo* screens.

Table S2 (relative to Figure 1). DUBs tested in the primary and secondary *in vivo* screens. Table S3 (relative to Figure 1). Results from *in vivo* RNAi screens.

Table S4. Primer sequences for Illumina sequencing multiplexing strategy. Table S5. Hematopoietic stem and progenitor cell populations and their markers.

Table S6. List of antibodies used in this study.

Table S7 (relative to Figure S4). Inflammation in *Usp15-/-* mice.

Table S8 (relative to Figures 6A, B and D). TCGA Pan-Cancer (PANCAN) samples.

## References

1. Adorno, M., Sikandar, S., Mitra, S.S., Kuo, A., Nicolis Di Robilant, B., Haro-Acosta, V., Ouadah, Y., Quarta, M., Rodriguez, J., Qian, D., et al. (2013). Usp16 contributes to somatic stem-cell defects in Down’s syndrome. Nature 501, 380–384.

2. Al-Salihi, M.A., Herhaus, L., Macartney, T., and Sapkota, G.P. (2012). USP11 augments TGFbeta signalling by deubiquitylating ALK5. Open Biol 2, 120063.

3. Arrigoni, E., Del Re, M., Galimberti, S., Restante, G., Rofi, E., Crucitta, S., Barate, C., Petrini, M., Danesi, R., and Di Paolo, A. (2018). Concise Review: Chronic Myeloid Leukemia: Stem Cell Niche and Response to Pharmacologic Treatment. Stem Cells Transl Med 7, 305–314.

4. Bagger, F.O., Rapin, N., Theilgaard-Monch, K., Kaczkowski, B., Thoren, L.A., Jendholm, J., Winther, O., and Porse, B.T. (2013). HemaExplorer: a database of mRNA expression profiles in normal and malignant haematopoiesis. Nucleic Acids Res 41, D1034–1039.

5. Baker, R.T., Wang, X.W., Woollatt, E., White, J.A., and Sutherland, G.R. (1999). Identification, functional characterization, and chromosomal localization of USP15, a novel human ubiquitin-specific protease related to the UNP oncoprotein, and a systematic nomenclature for human ubiquitin-specific proteases. Genomics 59, 264–274.

6. Bakker, S.T., and Passegue, E. (2013). Resilient and resourceful: genome maintenance strategies in hematopoietic stem cells. Exp Hematol 41, 915–923.

7. Beerman, I., Maloney, W.J., Weissmann, I.L., and Rossi, D.J. (2010). Stem cells and the aging hematopoietic system. Curr Opin Immunol 22, 500–506.

8. Biechonski, S., Yassin, M., and Milyavsky, M. (2017). DNA-damage response in hematopoietic stem cells: an evolutionary trade-off between blood regeneration and leukemia suppression. Carcinogenesis 38, 367–377.

9. Blank, U., and Karlsson, S. (2015). TGF-beta signaling in the control of hematopoietic stem cells. Blood 125, 3542–3550.

10. Cabezas-Wallscheid, N., Klimmeck, D., Hansson, J., Lipka, D.B., Reyes, A., Wang, Q., Weichenhan, D., Lier, A., von Paleske, L., Renders, S., et al. (2014). Identification of regulatory networks in HSCs and their immediate progeny via integrated proteome, transcriptome, and DNA Methylome analysis. Cell Stem Cell 15, 507–522.

11. Cellot, S., Hope, K.J., Chagraoui, J., Sauvageau, M., Deneault, E., MacRae, T., Mayotte, N., Wilhelm, B.T., Landry, J.R., Ting, S.B., et al. (2013). RNAi screen identifies Jarid1b as a major regulator of mouse HSC activity. Blood 122, 1545–1555.

12. Christensen, J.L., and Weissman, I.L. (2001). Flk-2 is a marker in hematopoietic stem cell differentiation: a simple method to isolate long-term stem cells. Proc Natl Acad Sci U S A 98, 14541–14546.

13. Citterio, E. (2015). Fine-tuning the ubiquitin code at DNA double-strand breaks: deubiquitinating enzymes at work. Front Genet 6, 282.

14. Clague, M.J., Coulson, J.M., and Urbe, S. (2012). Cellular functions of the DUBs. J Cell Sci 125, 277–286.

15. Das, T., Park, J.K., Park, J., Kim, E., Rape, M., Kim, E.E., and Song, E.J. (2017). USP15 regulates dynamic protein-protein interactions of the spliceosome through deubiquitination of PRP31. Nucleic Acids Res 45, 5010–5011.

16. de Haan, G., and Lazare, S.S. (2018). Aging of hematopoietic stem cells. Blood 131, 479–487.

17. Dey, A., Seshasayee, D., Noubade, R., French, D.M., Liu, J., Chaurushiya, M.S., Kirkpatrick, D.S., Pham, V.C., Lill, J.R., Bakalarski, C.E., et al. (2012). Loss of the Tumor Suppressor BAP1 Causes Myeloid Transformation. Science (New York, NY) 337, 1541–1546.

18. Doench, J.G., Fusi, N., Sullender, M., Hegde, M., Vaimberg, E.W., Donovan, K.F., Smith, I., Tothova, Z., Wilen, C., Orchard, R., et al. (2016). Optimized sgRNA design to maximize activity and minimize off-target effects of CRISPR-Cas9. Nat Biotechnol 34, 184–191.

19. Edgar, R., Domrachev, M., and Lash, A.E. (2002). Gene Expression Omnibus: NCBI gene expression and hybridization array data repository. Nucleic Acids Res 30, 207–210.

20. Eichhorn, P.J., Rodon, L., Gonzalez-Junca, A., Dirac, A., Gili, M., Martinez-Saez, E., Aura, C., Barba, I., Peg, V., Prat, A., et al. (2012). USP15 stabilizes TGF-beta receptor I and promotes oncogenesis through the activation of TGF-beta signaling in glioblastoma. Nat Med 18, 429–435.

21. Fielding, A.B., Concannon, M., Darling, S., Rusilowicz-Jones, E.V., Sacco, J.J., Prior, I.A., Clague, M.J., Urbe, S., and Coulson, J.M. (2018). The deubiquitylase USP15 regulates topoisomerase II alpha to maintain genome integrity. Oncogene.

22. Flach, J., Bakker, S.T., Mohrin, M., Conroy, P.C., Pietras, E.M., Reynaud, D., Alvarez, S., Diolaiti, M.E., Ugarte, F., Forsberg, E.C., et al. (2014). Replication stress is a potent driver of functional decline in ageing haematopoietic stem cells. Nature 512, 198–202.

23. Galeev, R., Baudet, A., Kumar, P., Rundberg Nilsson, A., Nilsson, B., Soneji, S., Torngren, T., Borg, A., Kvist, A., and Larsson, J. (2016). Genome-wide RNAi Screen Identifies Cohesin Genes as Modifiers of Renewal and Differentiation in Human HSCs. Cell Rep 14, 2988–3000.

24. Gargiulo, G., Serresi, M., Cesaroni, M., Hulsman, D., and van Lohuizen, M. (2014). In vivo shRNA screens in solid tumors. Nat Protoc 9, 2880–2902.

25. Gu, Y., Jones, A.E., Yang, W., Liu, S., Dai, Q., Liu, Y., Swindle, C.S., Zhou, D., Zhang, Z., Ryan, T.M., et al. (2016). The histone H2A deubiquitinase Usp16 regulates hematopoiesis and hematopoietic stem cell function. Proc Natl Acad Sci U S A 113, E51–60.

26. Haahr, P., Borgermann, N., Guo, X., Typas, D., Achuthankutty, D., Hoffmann, S., Shearer, R., Sixma, T.K., and Mailand, N. (2018). ZUFSP Deubiquitylates K63-Linked Polyubiquitin Chains to Promote Genome Stability. Mol Cell 70, 165–174 e166.

27. Harrigan, J.A., Jacq, X., Martin, N.M., and Jackson, S.P. (2018). Deubiquitylating enzymes and drug discovery: emerging opportunities. Nat Rev Drug Discov 17, 57–78.

28. Heideker, J., and Wertz, I.E. (2015). DUBs, the regulation of cell identity and disease. Biochem J 465, 1–26.

29. Heng, T.S., Painter, M.W., and Immunological Genome Project, C. (2008). The Immunological Genome Project: networks of gene expression in immune cells. Nat Immunol 9, 1091–1094.

30. Higuchi, M., Kawamura, H., Matsuki, H., Hara, T., Takahashi, M., Saito, S., Saito, K., Jiang, S., Naito, M., Kiyonari, H., et al. (2016). USP10 Is an Essential Deubiquitinase for Hematopoiesis and Inhibits Apoptosis of Long-Term Hematopoietic Stem Cells. Stem Cell Reports 7, 1116–1129.

31. Hope, K.J., Cellot, S., Ting, S.B., MacRae, T., Mayotte, N., Iscove, N.N., and Sauvageau, G. (2010). An RNAi screen identifies Msi2 and Prox1 as having opposite roles in the regulation of hematopoietic stem cell activity. Cell Stem Cell 7, 101–113.

32. Inui, M., Manfrin, A., Mamidi, A., Martello, G., Morsut, L., Soligo, S., Enzo, E., Moro, S., Polo, S., Dupont, S., et al. (2011). USP15 is a deubiquitylating enzyme for receptor-activated SMADs. Nat Cell Biol 13, 1368–1375.

33. Iyengar, P.V., Jaynes, P., Rodon, L., Lama, D., Law, K.P., Lim, Y.P., Verma, C., Seoane, J., and Eichhorn, P.J. (2015). USP15 regulates SMURF2 kinetics through C-lobe mediated deubiquitination. Sci Rep 5, 14733.

34. Jackson, S.P., and Durocher, D. (2013). Regulation of DNA damage responses by ubiquitin and SUMO. Mol Cell 49, 795–807.

35. Kiel, M.J., Yilmaz, Ö.H., Iwashita, T., Yilmaz, O.H., Terhorst, C., and Morrison, S.J. (2005). SLAM Family Receptors Distinguish Hematopoietic Stem and Progenitor Cells and Reveal Endothelial Niches for Stem Cells. Cell 121, 1109–1121.

36. Kreso, A., and Dick, J.E. (2014). Evolution of the cancer stem cell model. Cell Stem Cell 14, 275–291.

37. Kwasna, D., Abdul Rehman, S.A., Natarajan, J., Matthews, S., Madden, R., De Cesare, V., Weidlich, S., Virdee, S., Ahel, I., Gibbs-Seymour, I., et al. (2018). Discovery and Characterization of ZUFSP/ZUP1, a Distinct Deubiquitinase Class Important for Genome Stability. Mol Cell 70, 150–164 e156.

38. Lancini, C., Gargiulo, G., van den Berk, P.C., and Citterio, E. (2016). Quantitative analysis by next generation sequencing of hematopoietic stem and progenitor cells (LSK) and of splenic B cells transcriptomes from wild-type and Usp3-knockout mice. Data Brief 6, 556–561.

39. Lancini, C., van den Berk, P.C., Vissers, J.H., Gargiulo, G., Song, J.Y., Hulsman, D., Serresi, M., Tanger, E., Blom, M., Vens, C., et al. (2014). Tight regulation of ubiquitin-mediated DNA damage response by USP3 preserves the functional integrity of hematopoietic stem cells. J Exp Med 211, 1759–1777.

40. Laurenti, E., and Gottgens, B. (2018). From haematopoietic stem cells to complex differentiation landscapes. Nature 553, 418–426.

41. Liu, W.T., Huang, K.Y., Lu, M.C., Huang, H.L., Chen, C.Y., Cheng, Y.L., Yu, H.C., Liu, S.Q., Lai, N.S., and Huang, H.B. (2017). TGF-beta upregulates the translation of USP15 via the PI3K/AKT pathway to promote p53 stability. Oncogene 36, 2715–2723.

42. Long, L., Thelen, J.P., Furgason, M., Haj-Yahya, M., Brik, A., Cheng, D., Peng, J., and Yao, T. (2014). The U4/U6 recycling factor SART3 has histone chaperone activity and associates with USP15 to regulate H2B deubiquitination. J Biol Chem 289, 8916–8930.

43. McFarland, J.M., Ho, Z.V., Kugener, G., Dempster, J.M., Montgomery, P.G., Bryan, J.G., Krill-Burger, J.M., Green, T.M., Vazquez, F., Boehm, J.S., et al. (2018). Improved estimation of cancer dependencies from large-scale RNAi screens using model-based normalization and data integration. Nat Commun 9, 4610.

44. Mevissen, T.E.T., and Komander, D. (2017). Mechanisms of Deubiquitinase Specificity and Regulation. Annu Rev Biochem 86, 159–192.

45. Miller, P.G., Al-Shahrour, F., Hartwell, K.A., Chu, L.P., Jaras, M., Puram, R.V., Puissant, A., Callahan, K.P., Ashton, J., McConkey, M.E., et al. (2013). In Vivo RNAi screening identifies a leukemia-specific dependence on integrin beta 3 signaling. Cancer Cell 24, 45–58.

46. Morrison, S.J., and Spradling, A.C. (2008). Stem cells and niches: mechanisms that promote stem cell maintenance throughout life. Cell 132, 598–611.

47. Mu, J.J., Wang, Y., Luo, H., Leng, M., Zhang, J., Yang, T., Besusso, D., Jung, S.Y., and Qin, J. (2007). A proteomic analysis of ataxia telangiectasia-mutated (ATM)/ATM-Rad3-related (ATR) substrates identifies the ubiquitin-proteasome system as a regulator for DNA damage checkpoints. J Biol Chem 282, 17330–17334.

48. Naik, S.H., Perie, L., Swart, E., Gerlach, C., van Rooij, N., de Boer, R.J., and Schumacher, T.N. (2013). Diverse and heritable lineage imprinting of early haematopoietic progenitors. Nature 496, 229–232.

49. Nakagawa, M.M., Thummar, K., Mandelbaum, J., Pasqualucci, L., and Rathinam, C.V. (2015). Lack of the ubiquitin-editing enzyme A20 results in loss of hematopoietic stem cell quiescence. J Exp Med 212, 203–216.

50. Navarro, S., Meza, N.W., Quintana-Bustamante, O., Casado, J.A., Jacome, A., McAllister, K., Puerto, S., Surralles, J., Segovia, J.C., and Bueren, J.A. (2006). Hematopoietic dysfunction in a mouse model for Fanconi anemia group D1. Mol Ther 14, 525–535.

51. Nestorowa, S., Hamey, F.K., Pijuan Sala, B., Diamanti, E., Shepherd, M., Laurenti, E., Wilson, N.K., Kent, D.G., and Gottgens, B. (2016). A single-cell resolution map of mouse hematopoietic stem and progenitor cell differentiation. Blood 128, e20–31.

52. Nijnik, A., Clare, S., Hale, C., Raisen, C., Mcintyre, R.E., Yusa, K., Everitt, A.R., Mottram, L., Podrini, C., Lucas, M., et al. (2012). The critical role of histone H2A-deubiquitinase Mysm1 in hematopoiesis and lymphocyte differentiation. Blood 119, 1370–1379.

53. Nishi, R., Wijnhoven, P., le Sage, C., Tjeertes, J., Galanty, Y., Forment, J.V., Clague, M.J., Urbe, S., and Jackson, S.P. (2014). Systematic characterization of deubiquitylating enzymes for roles in maintaining genome integrity. Nat Cell Biol 16, 1016-1026, 1011–1018.

54. Oguro, H., Ding, L., and Morrison, S.J. (2013). SLAM family markers resolve functionally distinct subpopulations of hematopoietic stem cells and multipotent progenitors. Cell Stem Cell 13, 102–116.

55. Olsson, A., Venkatasubramanian, M., Chaudhri, V.K., Aronow, B.J., Salomonis, N., Singh, H., and Grimes, H.L. (2016). Single-cell analysis of mixed-lineage states leading to a binary cell fate choice. Nature 537, 698–702.

56. Orthwein, A., Noordermeer, S.M., Wilson, M.D., Landry, S., Enchev, R.I., Sherker, A., Munro, M., Pinder, J., Salsman, J., Dellaire, G., et al. (2015). A mechanism for the suppression of homologous recombination in G1 cells. Nature 528, 422–426.

57. Padmanabhan, A., Candelaria, N., Wong, K.K., Nikolai, B.C., Lonard, D.M., O’Malley, B.W., and Richards, J.S. (2018). USP15-dependent lysosomal pathway controls p53-R175H turnover in ovarian cancer cells. Nat Commun 9, 1270.

58. Park, I.-K., Qian, D., Kiel, M., Becker, M.W., Pihalja, M., Weissman, I.L., Morrison, S.J., and Clarke, M.F. (2003). Bmi-1 is required for maintenance of adult self-renewing haematopoietic stem cells. Nature 423, 302–305.

59. Parmar, K., Kim, J., Sykes, S., Shimamura, A., Stuckert, P., Zhu, K., Hamilton, A., Deloach, M., Kutok, J., Akashi, K., et al. (2010). Hematopoietic Stem Cell Defects in Mice with Deficiency of Fancd2 or Usp1. STEM CELLS 28, 1186–1195.

60. Pauli, E.K., Chan, Y.K., Davis, M.E., Gableske, S., Wang, M.K., Feister, K.F., and Gack, M.U. (2014). The ubiquitin-specific protease USP15 promotes RIG-I-mediated antiviral signaling by deubiquitylating TRIM25. Sci Signal 7, ra3.

61. Pellin, D., Loperfido, M., Baricordi, C., S.L., W., Montepeloso, A., Weinberg, O.K., Biffi, A., Klein, A.M., and Biasco, L. (2019). A comprehensive single cell transcriptional landscape of human hematopoietic progenitors. Nat Commun 10.

62. Peng, Y., Liao, Q., Tan, W., Peng, C., Hu, Z., Chen, Y., Li, Z., Li, J., Zhen, B., Zhu, W., et al. (2019). The deubiquitylating enzyme USP15 regulates homologous recombination repair and cancer cell response to PARP inhibitors. Nat Commun 10, 1224.

63. Pilzecker, B., Buoninfante, O.A., van den Berk, P., Lancini, C., Song, J.Y., Citterio, E., and Jacobs, H. (2017). DNA damage tolerance in hematopoietic stem and progenitor cells in mice. Proc Natl Acad Sci U S A 114, E6875–E6883.

64. Pritchard, C.E.J., Kroese, L.J., and Huijbers, I.J. (2017). Direct Generation of Conditional Alleles Using CRISPR/Cas9 in Mouse Zygotes. Methods Mol Biol 1642, 21–35.

65. Puram, R.V., Kowalczyk, M.S., de Boer, C.G., Schneider, R.K., Miller, P.G., McConkey, M., Tothova, Z., Tejero, H., Heckl, D., Jaras, M., et al. (2016). Core Circadian Clock Genes Regulate Leukemia Stem Cells in AML. Cell 165, 303–316.

66. Ritchie, M.E., Phipson, B., Wu, D., Hu, Y., Law, C.W., Shi, W., and Smyth, G.K. (2015). limma powers differential expression analyses for RNA-sequencing and microarray studies. Nucleic Acids Res 43, e47.

67. Schepers, K., Campbell, T.B., and Passegue, E. (2015). Normal and leukemic stem cell niches: insights and therapeutic opportunities. Cell Stem Cell 16, 254–267.

68. Schubert, M., Klinger, B., Klunemann, M., Sieber, A., Uhlitz, F., Sauer, S., Garnett, M.J., Bluthgen, N., and Saez-Rodriguez, J. (2018). Perturbation-response genes reveal signaling footprints in cancer gene expression. Nat Commun 9, 20.

69. Schweitzer, K., Bozko, P.M., Dubiel, W., and Naumann, M. (2007). CSN controls NF-kappaB by deubiquitinylation of IkappaBalpha. EMBO J 26, 1532–1541.

70. Schwertman, P., Bekker-Jensen, S., and Mailand, N. (2016). Regulation of DNA double-strand break repair by ubiquitin and ubiquitin-like modifiers. Nat Rev Mol Cell Biol 17, 379–394.

71. Sertorio, M., Du, W., Amarachintha, S., Wilson, A., and Pang, Q. (2017). In Vivo RNAi Screen Unveils PPARgamma as a Regulator of Hematopoietic Stem Cell Homeostasis. Stem Cell Reports 8, 1242–1255.

72. Srihari, S., and Ragan, M.A. (2013). Systematic tracking of dysregulated modules identifies novel genes in cancer. Bioinformatics 29, 1553–1561.

73. Teyra, J., Singer, A.U., Schmitges, F.W., Jaynes, P., Kit Leng Lui, S., Polyak, M.J., Fodil, N., Krieger, J.R., Tong, J., Schwerdtfeger, C., et al. (2019). Structural and Functional Characterization of Ubiquitin Variant Inhibitors of USP15. Structure 27, 590–605 e595.

74. Torre, S., Polyak, M.J., Langlais, D., Fodil, N., Kennedy, J.M., Radovanovic, I., Berghout, J., Leiva-Torres, G.A., Krawczyk, C.M., Ilangumaran, S., et al. (2017). USP15 regulates type I interferon response and is required for pathogenesis of neuroinflammation. Nat Immunol 18, 54–63.

75. Vasanthakumar, A., Arnovitz, S., Marquez, R., Lepore, J., Rafidi, G., Asom, A., Weatherly, M., Davis, E.M., Neistadt, B., Duszynski, R., et al. (2016). Brca1 deficiency causes bone marrow failure and spontaneous hematologic malignancies in mice. Blood 127, 310–313.

76. Vlasschaert, C., Cook, D., Xia, X., and Gray, D.A. (2017). The evolution and functional diversification of the deubiquitinating enzyme superfamily. Genome Biol Evol.

77. Vlasschaert, C., Xia, X., Coulombe, J., and Gray, D.A. (2015). Evolution of the highly networked deubiquitinating enzymes USP4, USP 15, and USP11. BMC Evol Biol 15, 230.

78. Walter, D., Lier, A., Geiselhart, A., Thalheimer, F.B., Huntscha, S., Sobotta, M.C., Moehrle, B., Brocks, D., Bayindir, I., Kaschutnig, P., et al. (2015). Exit from dormancy provokes DNA-damage-induced attrition in haematopoietic stem cells. Nature 520, 549–552.

79. Wang, J., Sun, Q., Morita, Y., Jiang, H., Groß, A., Lechel, A., Hildner, K., Guachalla, L.M., Gompf, A., Hartmann, D., et al. (2012). A Differentiation Checkpoint Limits Hematopoietic Stem Cell Self-Renewal in Response to DNA Damage. Cell 148, 1001–1014.

80. Wang, T., Nandakumar, V., Jiang, X.X., Jones, L., Yang, A.G., Huang, X.F., and Chen, S.Y. (2013). The control of hematopoietic stem cell maintenance, self-renewal, and differentiation by Mysm1-mediated epigenetic regulation. Blood 122, 2812–2822.

81. Warr, M.R., Pietras, E.M., and Passegue, E. (2011). Mechanisms controlling hematopoietic stem cell functions during normal hematopoiesis and hematological malignancies. Wiley Interdiscip Rev Syst Biol Med 3, 681–701.

82. Wijnhoven, P., Konietzny, R., Blackford, A.N., Travers, J., Kessler, B.M., Nishi, R., and Jackson, S.P. (2015). USP4 Auto-Deubiquitylation Promotes Homologous Recombination. Mol Cell 60, 362–373.

83. Wilson, A., Laurenti, E., Oser, G., van der Wath, R.C., Blanco-Bose, W., Jaworski, M., Offner, S., Dunant, C.F., Eshkind, L., Bockamp, E., et al. (2008). Hematopoietic stem cells reversibly switch from dormancy to self-renewal during homeostasis and repair. Cell 135, 1118–1129.

84. Yau, R., and Rape, M. (2016). The increasing complexity of the ubiquitin code. Nat Cell Biol 18, 579–586.

85. Ye, Z., Yu, X., and Cheng, L. (2008). Lentiviral gene transduction of mouse and human stem cells. Methods Mol Biol 430, 243–253.

86. Yeung, J., and Wai Eric So, A. (2009). Methods in Molecular Biology. 538, 301–315.

87. Zhang, L., Zhou, F., Drabsch, Y., Gao, R., Snaar-Jagalska, B.E., Mickanin, C., Huang, H., Sheppard, K.-A., Porter, J.A., Lu, C.X., et al. (2012). USP4 is regulated by AKT phosphorylation and directly deubiquitylates TGF-beta type I receptor. Nature Cell Biology 14, 1–12.

88. Zhang, X., Smits, A.H., van Tilburg, G.B., Jansen, P.W., Makowski, M.M., Ovaa, H., and Vermeulen, M. (2017). An Interaction Landscape of Ubiquitin Signaling. Mol Cell 65, 941–955 e948.

89. Zhou, F., Xie, F., Jin, K., Zhang, Z., Clerici, M., Gao, R., van Dinther, M., Sixma, T.K., Huang, H., Zhang, L., et al. (2017). USP4 inhibits SMAD4 monoubiquitination and promotes activin and BMP signaling. EMBO J 36, 1623–1639.

90. Zou, Q., Jin, J., Hu, H., Li, H.S., Romano, S., Xiao, Y., Nakaya, M., Zhou, X., Cheng, X., Yang, P., et al. (2014). USP15 stabilizes MDM2 to mediate cancer-cell survival and inhibit antitumor T cell responses. Nat Immunol 15, 562–570.

91. Zou, Q., Jin, J., Xiao, Y., Zhou, X., Hu, H., Cheng, X., Kazimi, N., Ullrich, S.E., and Sun, S.C. (2015). T Cell Intrinsic USP15 Deficiency Promotes Excessive IFN-gamma Production and an Immunosuppressive Tumor Microenvironment in MCA-Induced Fibrosarcoma. Cell Rep 13, 2470–2479.

92. Zuber, J., Shi, J., Wang, E., Rappaport, A.R., Herrmann, H., Sison, E.A., Magoon, D., Qi, J., Blatt, K., Wunderlich, M., et al. (2011). RNAi screen identifies Brd4 as a therapeutic target in acute myeloid leukaemia. Nature 478, 524–528.

