## Supplemental Information for "USP15 deubiquitinase safeguards hematopoiesis and genome integrity in hematopoietic stem cells and leukemia cells"

**Supplemental Figures**

Figure S1 (relative to Figure 1). *In vivo* RNAi screens for DUBs in murine hematopoietic stem and progenitor cells.
Figure S2 (relative to Figure 1). Pooled *in vivo* RNAi screens identify candidates DUBs effectors of HSC activity.
Figure S3 (relative to Figures 2C, 3, 4 and 5). Flow cytometry and gating strategies.
Figure S4. Generation and initial characterization of *Usp15* knockout mouse model (relative to Figure 4).
Figure S5. USP15 RNAi in CML cell lines results in alteration of inflammatory signaling pathways.
Figure S6. USP15 loss enhances genotoxic stress *in vitro* in human cancer cells and *in vivo* in mouse hematopoietic stem and progenitor cells.

**Supplemental Tables**

Table S1 (relative to Figure 1). Genes targeted and shRNAs used in the primary and secondary *in vivo* screens.
Table S2 (relative to Figure 1). DUBs tested in the primary and secondary *in vivo* screens.
Table S3 (relative to Figure 1). Results from *in vivo* RNAi screens.
Table S4. Primer sequences for Illumina sequencing multiplexing strategy.
Table S5. Hematopoietic stem and progenitor cell populations and their markers.
Table S6. List of antibodies used in this study.
Table S7 (Relative to Figure S4). Inflammation in *Usp15-/-* mice.
Table S8 (relative to Figures 6A, B and D). TCGA Pan-Cancer (PANCAN) samples used in the analysis.

**Figure S1 (relative to Figure 1). *In vivo* RNAi screens for DUBs in murine hematopoietic stem and progenitor cells.**

(A) Early progenitors (LSK) population is enriched and it is retained upon viral transduction *in vitro.* Lineage negative (Lin-) cells isolated form wt (CD45.2) bone marrow (BM) were grown under stem cell conditions and transduced with pooled shRNA lentiviral libraries targeting all annotated mouse orthologues of human DUBs. After selection, CD45.2 transduced Lin- cells are mixed 1:1 with CD45.1 support BM cells. Here, representative FACS profiles and relative percentage of LSK (Lin- Sca1+ cKit+) cells in freshly isolated Lin-, $cKit^{high}$ progenitors (left panel), in support BM (upper middle panel) and in shRNA library-transduced Lin-cells prior transplantation (upper right panel) is shown. Lower panels: LSK were gated on the basis of CD135/Flk-2 and SLAM CD150 surface markers to assess immature precursors endowed with reconstitution potential (LSK, CD135-, CD150+)(Christensen and Weissman, 2001; Kiel et al., 2005; Yeung and Wai Eric So, 2009). Frequency in the parental cell population is shown.

(B-C) ShRNA library-transduced Lin- cells efficiently engraft and contribute to BM repopulation (4 wpt). Representative FACS profiles (B) and quantification of engraftment for each screen (C) (primary screen, left panels; secondary screens, middle and right panels) is shown. Total reconstitution of peripheral blood cells (PBC) and contribution of donor-derived, transduced (CD45.2) Lin- cells to Gr1+, CD19+ (B cells) and CD3+ (T cells) in the blood (C, upper panels), BM (total BM cells, Lin-, cKit+ cells and LSK cells)(C, middle panels), and to splenic CD43- and B cells (CD43-, CD19+, CD220+)(C, bottom panels) in recipient mice at 4 wpt is shown. Error bars represent ± SEM.

For isolation of resting B cells from spleen single cell suspension, CD43 positive cells were first depleted using anti-mouse CD43 (Ly-48). Donor-derived cells were then phenotypically analyzed and isolated by FACS sorting using fluorochrome-labeled antibodies.

**Figure S2 (relative to Figure 1). Pooled *in vivo* RNAi screens identify candidates DUBs effectors of HSC activity.**

(A-D) Primary screen, full DUB library.

(A) shRNAs library complexity is maintained in transduced Lin- cells. Pairwise correlation between the shRNA-seq profiles of the plasmid DNA (plasmid library) used to generate the viral pool and the genomic DNA isolated from Lin- cells infected with the same viral pool prior transplantation, INPUT (T0).

(B) PC analysis of *in vivo* Lin- cells harvested from recipient mice at 4 wpt and INPUT (T0) samples.

(C) Quantitative heat map depiction of the primary screen with individual shRNAs (in rows) in each replicate (column) shown in red for the highest number of reads and in blue for lowest numbers of reads.

(D) Scatter plot of the change in shRNA representation after *in vivo* growth shows good correlation between the BM and the spleen compared to the input. Person's r is indicated.

(E) Secondary screen: DUB1 sub-library. Left panel: consolidated fraction of shRNAs retrieved *in vivo* in BM Lin- cells at 4 wpt and in controls. Right panel: PC analysis of *in vivo* and INPUT (T0) samples.

(F) Quantitative heat map depiction of the secondary screen (DUB2 sub-library) with individual shRNAs (in rows) in each replicate (column) shown in red for the highest number of reads and in blue for lowest numbers of reads.

(G) Secondary screen: DUB2 sub-library. Left panel: consolidated fraction of shRNAs retrieved *in vivo* in BM Lin- cells at 4 wpt and in controls. Right panel: PC analysis of *in vivo* and INPUT (T0) samples.

(H) Classification of screen results depicting significantly differentially represented DUBs overlapping between the primary and secondary screens. Candidate genes targeted by at least two shRNAs were selected with the indicated criteria. Genes were not scored if multiple cognate hairpins showed opposing effects. The number (#) of differentially represented shRNAs per gene is indicated.

(I-K) USP15 is highly and homogeneously expressed in the hematopoietic lineage tree at single cell level.

(I) scRNA-Seq on stem/multipotent progenitors (LSK; lin−Sca1+c-Kit+), common myeloid progenitors (CMP), granulocyte monocyte progenitors (GMP), and LK CD34+ cells (lin−c-Kit+CD34+) that included granulocytic precursors. HSCP-1, hematopoietic stem cell progenitor; HSCP-2; Meg, megakaryocytic; Eryth, erythrocytic; Multi-Lin*, multi-lineage primed; MDP, monocyte-dendritic cell precursor; Mono, monocytic; Gran, granulocytic; and myelocyte, myelocytes and metamyelocytes. EndFragment. Mouse Expression (Olsson et al., 2016).

(J) scRNA-seq on HSC/hematopoietic progenitors. LSK, hematopoietic stem cells and hematopoietic multipotent progenitors; LT-HSC, long-term hematopoietic stem cells (Lin–c-

Kit+Sca1+CD34–Flk2–); MEP-CMP-GMP, megakaryocyte-erythroid progenitor cells (MEP), common myeloid progenitors (CMP), and granulocyte-monocyte progenitors. Mouse Expression (Nestorowa et al., 2016).

(K) USP15 is expressed in single cells human CD34+ HSC and progenitors.

Analysis of USP15 expression in scRNA-seq data set of the human Lin−compartment, as investigated by means of CD34/CD164 fractionation (Pellin et al., 2019). The panels represent a SPRING plot of sorted human Lin−CD34/CD164 single-cell transcriptomes. Each point is one cell. Labels at the edges represent the transcriptional states associated to early lineage commitment (P, early progenitor cells; Meg, megakaryocytes; E, erythroid cells; BaP, basophilprogenitors;N, neutrophils; M, monocytes; DC, dendritic cells; Ly-T/B/NK, lymphoid T/B/NK cells) (Pellin et al., 2019). Plots of USP15 expression and of the lineage defining genes HLF (P, HSC) and ID2 (B-lymphocytes progenitors branch) (LRT adjusted pvalue $< 0.05$) is shown in each panel.

**Figure S3 (relative to Figures 2C, 3, 4 and 5). Flow cytometry and gating strategies.**

(A) Flow cytometry analysis of Lin- cells transduced with the indicated shRNA and grown *in vitro*. FACS representative profiles and LSK frequency per live cells are shown (relative to Figure 2C).

(B, C) Gating strategy for blood and spleen subsets analysis in bone marrow transplantation assays (relative to Figure 3 and 5).

(D) Gating strategy for hematopoietic stem and progenitor compartment analysis (relative to Figure 3, 4 and 5). Within the lineage-, Sca1+, c-Kit+ (LSK) population, cells were gated based on CD48 and CD150 to identify HSC (Cabezas-Wallscheid et al., 2014; Kiel et al., 2005).

Within the lineage-, c-Kit+, Sca1- (LKS-) population, cells were gated based on CD16/32 and CD34 expression to identify common myeloid progenitors (CMP), granulocyte monocyte progenitors (GMP) and megakaryocyte erythrocyte progenitors (MEP).

**Figure S4. Generation and initial characterization of *Usp15* knockout mouse model.**

(A) Schematic representation of the strategy for germline deletion of the entire mouse *Usp15* locus (91,795 bp) by CRISPR/Cas9. On top: wild-type *Usp15* genomic locus. Oligos (arrows) for PCR screening are indicated. Middle: targeting primers positioned at the transcriptional start site within exon 1 (5’ Target) and after the PolyA signal in exon 22 (3’ Target). Bottom: PCR fragments corresponding to the *Usp15* knockout (KO) or wild-type (wt) allele.

(B) PCR analysis of genomic tail tip DNA.

(C) Immunoblot of whole cell extract from freshly isolated BM cells form *Usp15+/+* (n=3) and *Usp15-/-* (n=3) mice with anti-USP15 antibody. Total proteins staining with Ponceau S is shown (lower panel).

(D) Genotype frequency per litter, on a total of 31 litters. n = number of born mice/genotype.

(E) Kaplan-Meier survival analysis.

(F) Body weight of mice of the indicated genotype. Males, 8-12 wks old. Females, 8-14 wks old.

(G) Histopathological analysis of *Usp15-/-* mice. Full pathological analysis was performed on 6 *Usp15+/+* and 5 *Usp15-/-* mice between 10 and 12 weeks of age. 3 out of 5 *Usp15-/-* animals showed inflammatory lesions, in one case severe (bottom). Top: representative H&E-stained head from a *Usp15-/-* mouse (11 wks) showing lesions of otitis media in one of the inner ears. Proteaceous material (*) with dead cells and inflammatory cells (black arrow) is visible, as well as edema (green arrow). Bottom: Representative H&E-stained intestine from a *Usp15-/-* mouse (11 wks) showing severe inflammatory lesions with massive immune cells infiltration in the mesentery (black arrows). No lesions were found in *Usp15+/+* littermates (not shown).

(H) Immunohistochemistry (IHC) of Ly6-G and F4/80 confirmed the nature of multiple inflammatory lesions encountered in the skin and abdominal cavity of the mouse (case n.1 in Table S7). As shown in the representative microphotographs of mesentery stained by H&E (Hematoxylin/Eosin) and IHC, the cells involved in the massive inflammatory infiltrations were mainly those neutrophils and macrophages that were labeled by Ly6-G and F4/80, respectively. The inflammation was accompanied by multiple necrotic changes. Thereby a nature of acute purulent inflammation is evident. Bar: 50 μm.

(I) Bone marrow was isolated from femurs of *Usp15+/+* (n=9) and *Usp15-/-* (n=5) 8-12 wks old mice. The number of total BM live cells was calculated and normalized to *Usp15+/+* littermates.

(J-R) Hematopoietic primitive compartment in USP15 knockout mice (relative to Figure 4).

(J-M) Multiparameter flow cytometry analysis of the hematopoietic primitive populations in *Usp15+/+* and *Usp15-/-* mice (8-12 wks old).

(J) Gating strategy with representative FACS profiles using CD135/Flk-2 and CD150 surface markers to assess immature precursors endowed with reconstitution potential (LSK, CD135-, CD150+)(Christensen and Weissman, 2001; Kiel et al., 2005; Yeung and Wai Eric So, 2009).

(K) LSK, CD150+, CD135- cell numbers per million live BM cells.

(L) Frequency of LSK, CD150+, CD135- immature precursors in bone marrow of *Usp15-/-* mice was calculated and normalized to *Usp15+/+* animals (*Usp15+/+* n=9, *Usp15-/- n=5*).

(M) BM frequency of the indicated progenitor populations in *Usp15-/-* mice normalized to Usp15+/+ animals. Results are from three (K, L, and M left panel) (*Usp15+/+* n=9, *Usp15-/- n=5*) or two (M, right panel) independent experiments.

(N) BM cells from wt or *Usp15*-/- mice were assayed for myeloid colony-forming (CFU-C) ability. Results are the mean ± SEM from two independent experiments, *n* = 3 per genotype per experiment. Representative images of two technical replica (A, B) are shown.

(O) Cell cycle analysis of hematopoietic stem and progenitor cells populations in wt and USP15 knockout mice. Percentage of G0/G1 and S/G2 cells in *Usp15+/+* and *Usp15-/-* mice was determined by DAPI analysis in 12-16 weeks old mice. *Usp15+/+*, n=10; *Usp15-/-*, n=5. Results are the mean ± SEM of three independent experiments.

(P) Cleaved (cl.) Caspase-3 analysis of freshly isolated hematopoietic stem and progenitor cells in *Usp15*+/+ and *Usp15*-/- mice. Results are the mean ± SEM of two independent experiments. *Usp15+/+*, n=6; *Usp15-/-*, n=2. 15-20 weeks old mice.

(Q) Representative images of *Usp15*+/+ and *Usp15-/-* BM sections stained for apoptosis-indicating cleavage (cl.) of Caspase-3. *n* = 6 mice per genotype. Bar, 500 μm.

(R) RNA-seq of LSK population freshly isolated from BM. Comparison of normalized gene expression data for *Usp15*-/- (n-=2) versus *Usp15*+/+ (n=3) LSK (2 months old mice). USP15 expression is highlighted in red.

For all panels: **, P ≤ 0.01; ***, P ≤ 0.001; ****, P ≤ 0.0001. P-value was assessed by Student's *t* test (D, F, K, L) or by Log-rank test (E). Mean ± SEM is shown.

**Figure S5. USP15 RNAi in CML cell lines results in alteration of inflammatory signaling pathways.**

(A) The dependency scores were calculated by Demeter2 for USP15 RNAi in CCLE lines (McFarland et al., 2018) and correlated to USP15 gene expression levels. TPM (Transcripts Per Million).

(B) Differential pathway activation by PROGENy for cell lines with DEMETER2 scores D2<-0.2 (Low) and D2>0.2 (high).

(C) Heat map showing PROGENy scores for the indicated pathways (rows) in all the CCLE cell lines (columns). Rows are sorted by correlation whereas columns are sorted by DEMETER2 score. The legends indicate DEMETER2 score and tumor type.

(D-I) KBM7 (D, E, F, G) or K562 (F, H, I) cells were transfected with USP15 (siUSP15) or non-targeting (siCtrl) siRNA pool. At 72 hours after transfection, cells were harvested and RNA was extracted and subjected to RNAseq (D, E, F, I) or used for RT-qPCR analysis (G, H).

(D) Unbiased clustering of genes significantly dysregulated (>1 fold change and Adjusted *P* value <0.05) in KBM7 RNAseq dataset.

(E) Ingenuity pathway analysis of USP15-dependent genes in the KBM7 dataset showing dysregulated top canonical pathways (cut-off logFC > 1; adjusted P value <0.05).

(F) Venn diagram depicting the overlap between Top 50 significantly enriched Ingenuity Upstream Regulators in each cell line RNAseq datasets. The Upstream Regulators were calculated based on differential gene expression between USP15 and control siRNAs. The nine overlapping terms are indicated below.

(G) qRT-PCR analysis of the indicated genes in KBM7 USP15 knock down cells.

(H) qRT-PCR analysis of the indicated genes in K562 USP15 knock down cells.

(I) USP15 RNAi in K562 cells. Ingenuity pathway analysis of USP15-dependent genes in the K562 RNAseq dataset showing dysregulated Top Regulator Effect Networks (cut-off logFC > 1; adjusted P value <0.05).

Results are from one (KBM7) or two independent experiments (K562) per cell line. Each experiment n=2 per siRNA/ cell line. Results are mean ± SD (F, G). **. $P \leq 0.01$; ***. $P \leq 0.001$; ****. $P \leq 0.0001$. P-value was assessed by Student's *t* test.

**Figure S6. USP15 loss enhances genotoxic stress *in vitro* in human cancer cells and *in vivo* in mouse hematopoietic stem and progenitor cells.**

(A-G) Knockdown or CRISPR/Cas9- mediated knockout of USP15 in U2OS cancer cells induces spontaneous DNA damage.

(A-F) U2OS cells were transfected with USP15 (siUSP15) or non-targeting (siCtrl) siRNA pool and assayed at 72 hours after transfection.

(A) Representative images of co-immunostaining for γH2AX and 53BP1. Arrows indicate micronuclei (MN), nucleoplasmatic bridges (NPB), and abnormal mitosis.

(B) Western blot with the indicated antibodies on whole cell extract.

(C) Quantification by Image J of the number of spontaneous 53BP1 foci per cell. n=4 experiments; >400 cells per genotype were in total analyzed. Mean values ± SEM are shown.

(D) Quantification of the percentage of cells containing >5 spontaneous γH2AX foci. Mean ± SD is shown.

(E) Percentage of micronuclei. Mean values ± SEM are shown.

(F) Percentage of aberrant mitosis, including bi-nucleated cells and cells with nucleoplasmatic bridges. Mean values ± SEM are shown.

(A-F) All quantifications are form three independent experiments. *n* > 50 cells/genotype/experiment.

(G-H) CRISPR/Cas9- mediated deletion of USP15 in U2OS cells.

(G) Western blot with the indicated antibodies on whole cell extract.

(H) Micronuclei were quantified in wt cells and in two independent USP15 knockout clones, in two independent experiments. A minimum of 100 cells per genotype/clone/experiment were scored. Mean values ± SEM are shown.

(I-K) USP15 protects the hematopoietic stem and progenitor compartment from genotoxic stress *in vivo*. *Usp15+/+ and Usp15-/-* mice (8-12 wks old) were injected with PBS or with low dose of cisplatin (CsPt)(0.8 mg/kg i.v.). Mice were sacrificed after 48 hrs.

(I) BM cells were isolated from wt or *Usp15*-/- mice and were assayed for myeloid colony-forming (CFU-C) ability.

(J, K) Multiparameter flow cytometry analysis of the hematopoietic stem and progenitor compartment.

(J) HSC (LSK, CD150+, CD48-) cell numbers per million live BM cells.

(K) The frequency of the indicated cell populations in *Usp15-/-* mice was calculated and normalized to *Usp15+/+* animals. LSK/CD150+ CD48- (HSC), lineage-, c-Kit+, Sca1- (LKS-), common myeloid progenitors (CMP), granulocyte monocyte progenitors (GMP), megakaryocyte erythrocyte progenitors (MEP), common lymphoid progenitors (CLP).

Results are the mean ± SEM or SD (K, left panel) of two independent experiments. Mice: (J-K) *Usp15+/+* $n = 4$, *Usp15-/-* $n = 3$ per treatment; (K) $n = 3$ per genotype per treatment.

(L) Kasumi-1 and MV4-11 cells harboring USP15 shRNA were kept in medium with or without doxycycline (Dox) for 5 days. Western blot was performed with the indicated antibodies. One representative experiment out of 2 is shown.

For all panels: *, $P \leq 0.05$; **,$P \leq 0.01$; ***, $P \leq 0.001$; ****, $P \leq 0.001$ as assessed by Student's t test.

### Supplemental Tables

**Table S1. Genes targeted and shRNAs used in the primary and secondary *in vivo* screens**
**Table S2. DUBs tested in the primary and secondary *in vivo* screens**
**Table S3. Results from *in vivo* RNAi screens**
Tables S1, S2 and S3 (relative to Figure 1) are included as separate Excel files.

**Table S4. Primer sequences for Illumina sequencing multiplexing strategy**

| | |
|---|---|
| **PCR 1, forward primer** | *ACACTCTTTCCCTACACGACGCTCTTCCGATCT*NNNNNNGGCTTTATATATCTTGTGGAAAGGACG |
| **PCR 1, reverse primer** | GTGACTGGAGTTCAGACGTGTGCTCTTCCGATCTGTGGATGAATACTGCCATTTGTCTC |
| **PCR 2, forward primer** | AATGATACGGCGACCACCGAGATCT*ACACTCTTTCCCTACACGACGCTCTTCCGATCT* |
| **PCR 2, reverse primer** | CAAGCAGAAGACGGCATACGAGAT**NNNNNN**GTGACTGGAGTTCAGACGTGTGCTCTTCCGATCT |
| **PCR 1 barcodes (green)** | |
| Barcode #1 | CGTGAT |
| Barcode #2 | ACATCG |
| Barcode #3 | GCCTAA |
| Barcode #4 | TGGTCA |
| Barcode #5 | CACTGT |
| Barcode #6 | ATTGGC |
| Barcode #7 | GATCTG |
| Barcode #8 | TCAAGT |
| Barcode #9 | CTGATC |
| Barcode #10 | AAGCTA |
| Barcode #11 | GTAGCC |
| Barcode #12 | TACAAG |
| **PCR 2 Illumina Indexes (green)** | |
| Index #1 | ATCACG |
| Index #2 | CGATGT |
| Index #3 | TTAGGC |
| Index #4 | TGACCA |

PCR1 forward primer is designed to include: Illumina/Solexa adapter 1 (red, italics), barcode (green, bold and underlined), pLKO.1 common region (plain text).

**Table S5. Hematopoietic stem and progenitor cell populations and their markers**

| Subset | Subpopulations | Markers |
|---|---|---|
| $Lin^-$ | LSK, HSC, LKS-, CMP, GMP, MEP | $Lineage^-$ |
| LSK | HSC | $Lineage^-$, $Sca\text{-}1^+$, $cKit^+$ |
| HSC | | $Lineage^-$, $Sca\text{-}1^+$, $cKit^+$, $CD150^+$, $CD48^-$ |
| $LKS^-$ | CMP, GMP, MEP | $Lineage^-$, $Sca\text{-}1^-$, $cKit^+$ |
| CMP | | $Lineage^-$, $Sca\text{-}1^-$, $cKit^+$, $CD34^{int}$, $CD16/32^{int}$ |
| GMP | | $Lineage^-$, $Sca\text{-}1^-$, $cKit^+$, $CD34^+$, $CD16/32^+$ |
| MEP | | $Lineage^-$, $Sca\text{-}1^-$, $cKit^+$, $CD34^-$, $CD16/32^-$ |

**Table S6. List of antibodies used in this study**

| Antigen-conjugate | Supplier | Dilution |
|---|---|---|
| Lineage Cell Detection Mixture-Biotin, mouse | MACS Miltenyi Biotech | 1:30 |
| Streptavidin APC/Cy7 | Southern Biotech | 1:200 |
| cKit-APC | eBioscience | 1:200 |
| Sca1-PerCp-Cy5.5 | Biolegend | 1:200 |
| CD48-FITC | BD Biosciences | 1:200 |
| CD135-PE | Biolegend | 1:200 |
| CD150-PE-Cy7 | Biolegend | 1:200 |
| CD16/32-PE-Cy7 | eBioscience | 1:200 |
| CD34-FITC | eBioscience | 1:200 |
| CD45.1(Ly5.1)-EF450 | eBioscience | 1:200 |
| CD45.2(Ly5.2)-PE | Biolegend | 1:200 |
| CD45.1(Ly5.1)-PE | BD Biosciences | 1:200 |
| CD45.2(Ly5.2)-PE-Cy7 | Biolegend | 1:200 |
| CD3-FITC | Biolegend | 1:200 |
| CD19-APC | BD Biosciences | 1:200 |
| Gr1-APC-Cy7 | Biolegend | 1:200 |
| CD11b-PerCp-Cy5.5 | BD Biosciences | 1:200 |
| CD4-APC | Biolegend | 1:200 |
| CD8a-PerCp-C5.5 | BD Biosciences | 1:200 |
| CD19-APC-H7 | BD Biosciences | 1:200 |
| CD43 (Ly-48) MicroBeads, mouse | MACS Miltenyi Biotech | n.a. |
| CD43-biotin (clone S7, RUO) | BD Biosciences | 1:100 |
| CD19-APC | Biolegend | 1:200 |
| B220-Pacific Blue | BD Biosciences | 1:200 |
| CD45.2-FITC | BD Biosciences | 1:200 |
| γH2AX | Millipore | 1:1000 |
| 53BP1 | Novus Biologicals | 1:500 |
| USP15 | Abcam | 1:500 |
| Tubulin | Sigma | 1:10,000 |
| Goat anti-mouse HRP | Life technologies | 1:10,000 |
| Alexa Fluor 568 goat anti–mouse | Life Technologies | 1:250 |
| Alexa Fluor 488 goat anti–rabbit | Life Technologies | 1:250 |

**Table S7 (Relative to Figure S4). Inflammation in *Usp15-/-* mice**

| Genotype | Age | Cases | Sign of inflammation |
|---|---|---|---|
| *Usp15+/+* | 10-12 wks | 0/6 | No lesions detected |
| *Usp15-/-* | 10-12 wks | 3/5 | *Case 1:*<br>Peritonitis, Laryngitis.<br>Skin: Local purulent inflammatory lesions subcutaneously, showing massive infiltration of neutrophils accompanied by necrosis of the tissue.<br>Testicular: degeneration in parts of the testes, accompanied by severe inflammatory changes in the surrounding adipose tissues with adiponecrosis.<br>GI tract: Purulent inflammatory lesions in the mesentery accompanied by fibrosis and adherence.<br>Spleen: Enhancement of haematopoiesis in the red pulp.<br>Pancreas: Edematous changes with inflammatory infiltrations in stroma.<br>Kidney: Purulent inflammatory lesions in the renal capsules and in the surrounding adipose tissues.<br>Gall bladder: Edematous changes in the submucosa with inflammatory lesions in the surroundings.<br>Head: Edema and inflammatory infiltrations in the submucosa of larynx.<br>Spleen: Enhancement of hematopoiesis in the red pulp of spleen<br><br>*Case 2:*<br>Head: -Local purulent inflammatory lesions in the skin of lip.<br>- Otitis media in one of the middle ears.<br>-Uterus/ovary Edematous changes of the endometrial stroma.<br><br>*Case 3:*<br>Head: Lesions of otitis media in one of the middle ears. |

Histopathological analysis of *Usp15-/-* mice. Full pathological analysis was performed on 6 *Usp15+/+* and 5 *Usp15-/-* mice between 10 and 12 weeks (wks) of age. GI: gastro intestinal tract.

**Table S8 (relative to Figures 6A, B and D). TCGA Pan-Cancer (PANCAN) samples**

| PANCAN EXPRESSION analysis (A,B) | | | PANCAN survival analysis (C) | |
|---|---|---|---|---|
| acute myeloid leukemia | 173 | | acute myeloid leukemia | 161 |
| adrenocortical cancer | 79 | | adrenocortical cancer | 79 |
| bladder urothelial carcinoma | 427 | | bladder urothelial carcinoma | 426 |
| brain lower grade glioma | 530 | | brain lower grade glioma | 528 |
| breast invasive carcinoma | 1218 | | breast invasive carcinoma | 1214 |
| cervical & endocervical cancer | 310 | | cervical & endocervical cancer | 309 |
| cholangiocarcinoma | 45 | | cholangiocarcinoma | 45 |
| colon adenocarcinoma | 495 | | colon adenocarcinoma | 491 |
| diffuse large B-cell lymphoma | 48 | | diffuse large B-cell lymphoma | 48 |
| esophageal carcinoma | 196 | | esophageal carcinoma | 196 |
| glioblastoma multiforme | 172 | | glioblastoma multiforme | 166 |
| head & neck squamous cell carcinoma | 566 | | head & neck squamous cell carcinoma | 565 |
| kidney chromophobe | 91 | | kidney chromophobe | 89 |
| kidney clear cell carcinoma | 606 | | kidney clear cell carcinoma | 606 |
| kidney papillary cell carcinoma | 323 | | kidney papillary cell carcinoma | 322 |
| liver hepatocellular carcinoma | 423 | | liver hepatocellular carcinoma | 422 |
| lung adenocarcinoma | 576 | | lung adenocarcinoma | 567 |
| lung squamous cell carcinoma | 553 | | lung squamous cell carcinoma | 546 |
| mesothelioma | 87 | | mesothelioma | 86 |
| ovarian serous cystadenocarcinoma | 308 | | ovarian serous cystadenocarcinoma | 307 |
| pancreatic adenocarcinoma | 183 | | pancreatic adenocarcinoma | 183 |
| pheochromocytoma & paraganglioma | 187 | | pheochromocytoma & paraganglioma | 187 |
| prostate adenocarcinoma | 550 | | prostate adenocarcinoma | 550 |
| rectum adenocarcinoma | 171 | | rectum adenocarcinoma | 170 |
| sarcoma | 265 | | sarcoma | 265 |
| skin cutaneous melanoma | 474 | | skin cutaneous melanoma | 458 |
| stomach adenocarcinoma | 450 | | stomach adenocarcinoma | 443 |
| testicular germ cell tumor | 156 | | testicular germ cell tumor | 139 |
| thymoma | 122 | | thymoma | 121 |
| thyroid carcinoma | 572 | | thyroid carcinoma | 572 |
| uterine carcinosarcoma | 57 | | uterine carcinosarcoma | 57 |
| uterine corpus endometrioid carcinoma | 567 | | uterine corpus endometrioid carcinoma | 553 |
| uveal melanoma | 80 | | uveal melanoma | 80 |
| Total number of samples | 11060 | | | 10951 |

### References

Cabezas-Wallscheid, N., Klimmeck, D., Hansson, J., Lipka, D.B., Reyes, A., Wang, Q., Weichenhan, D., Lier, A., von Paleske, L., Renders, S., *et al.* (2014). Identification of regulatory networks in HSCs and their immediate progeny via integrated proteome, transcriptome, and DNA Methylome analysis. Cell Stem Cell *15*, 507-522.

Christensen, J.L., and Weissman, I.L. (2001). Flk-2 is a marker in hematopoietic stem cell differentiation: a simple method to isolate long-term stem cells. Proc Natl Acad Sci U S A *98*, 14541-14546.

Kiel, M.J., Yilmaz, Ö.H., Iwashita, T., Yilmaz, O.H., Terhorst, C., and Morrison, S.J. (2005). SLAM Family Receptors Distinguish Hematopoietic Stem and Progenitor Cells and Reveal Endothelial Niches for Stem Cells. Cell *121*, 1109-1121.

McFarland, J.M., Ho, Z.V., Kugener, G., Dempster, J.M., Montgomery, P.G., Bryan, J.G., Krill-Burger, J.M., Green, T.M., Vazquez, F., Boehm, J.S., *et al.* (2018). Improved estimation of cancer dependencies from large-scale RNAi screens using model-based normalization and data integration. Nat Commun *9*, 4610.

Nestorowa, S., Hamey, F.K., Pijuan Sala, B., Diamanti, E., Shepherd, M., Laurenti, E., Wilson, N.K., Kent, D.G., and Gottgens, B. (2016). A single-cell resolution map of mouse hematopoietic stem and progenitor cell differentiation. Blood *128*, e20-31.

Olsson, A., Venkatasubramanian, M., Chaudhri, V.K., Aronow, B.J., Salomonis, N., Singh, H., and Grimes, H.L. (2016). Single-cell analysis of mixed-lineage states leading to a binary cell fate choice. Nature *537*, 698-702.

Pellin, D., Loperfido, M., Baricordi, C., S.L., W., Montepeloso, A., Weinberg, O.K., Biffi, A., Klein, A.M., and Biasco, L. (2019). A comprehensive single cell transcriptional landscape of human hematopoietic progenitors. Nat Commun *10*.
Yeung, J., and Wai Eric So, A. (2009). Methods in Molecular Biology. *538*, 301-315.

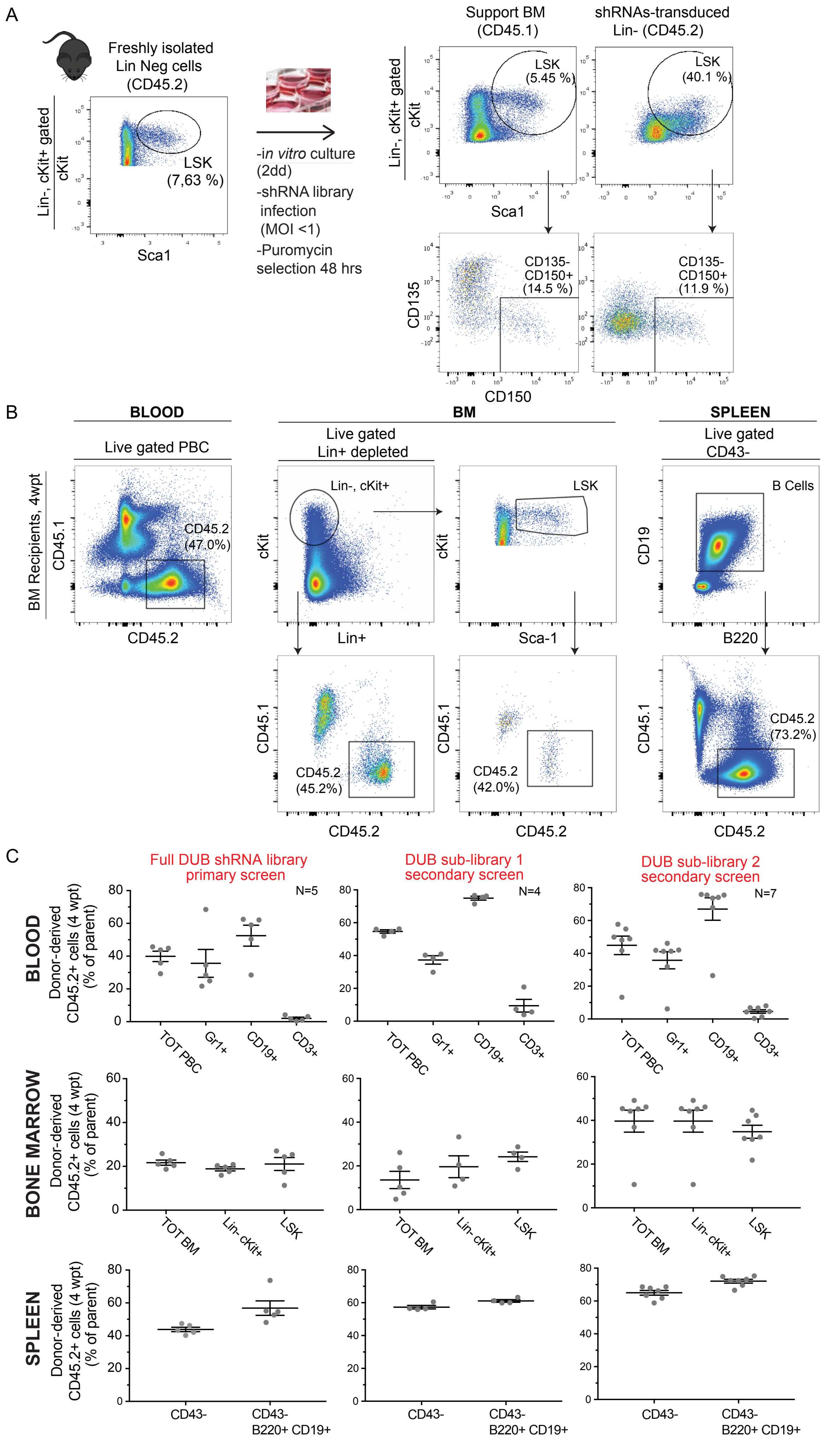

**Figure S1.** ***In vivo*** **RNAi screens for DUBs in murine hematopoietic stem and progenitor cells (relative to Figure 1).**

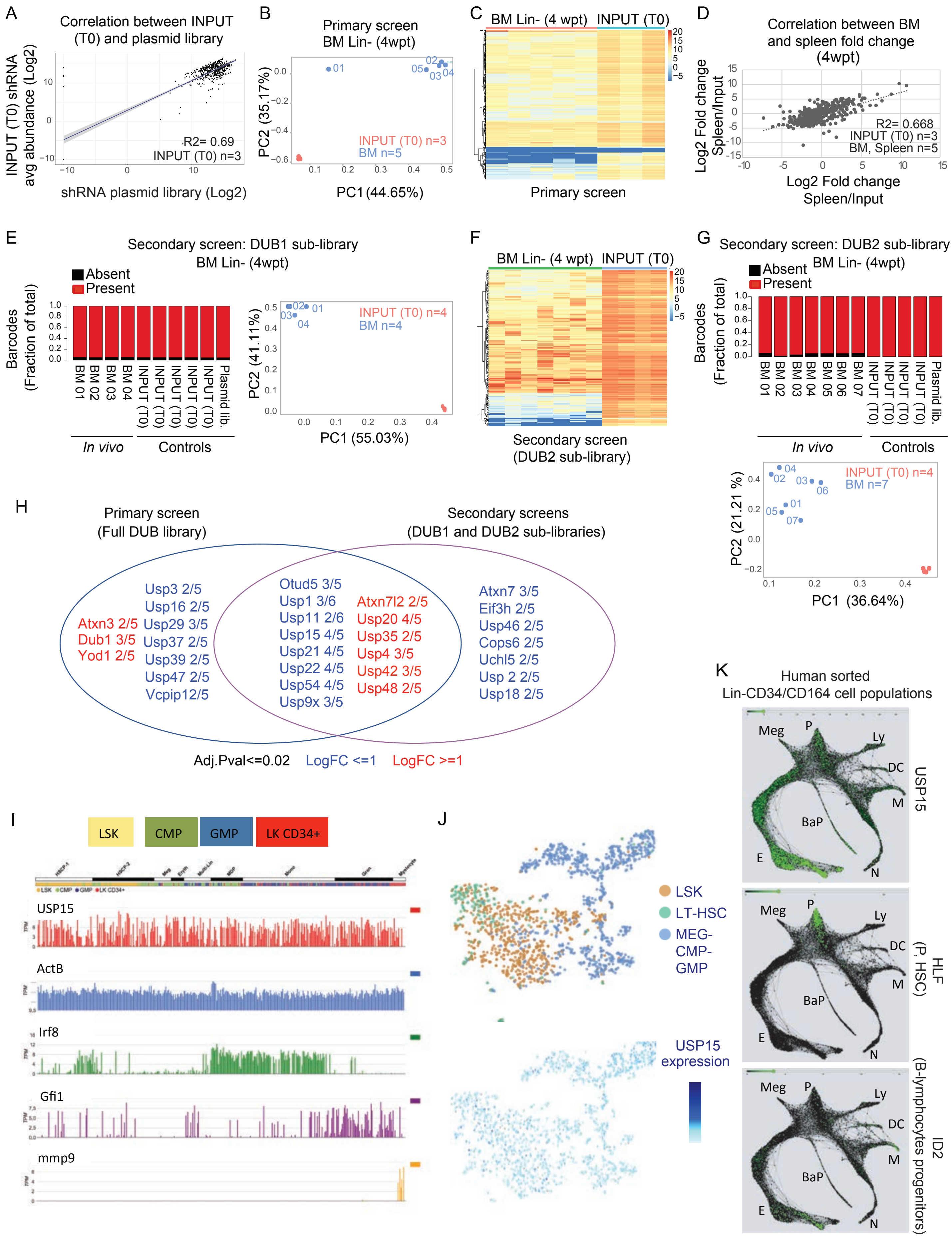

**Figure S2. Pooled *in vivo* RNAi screens identify candidate DUBs effectors of HSC activity (relative to Figure 1).**

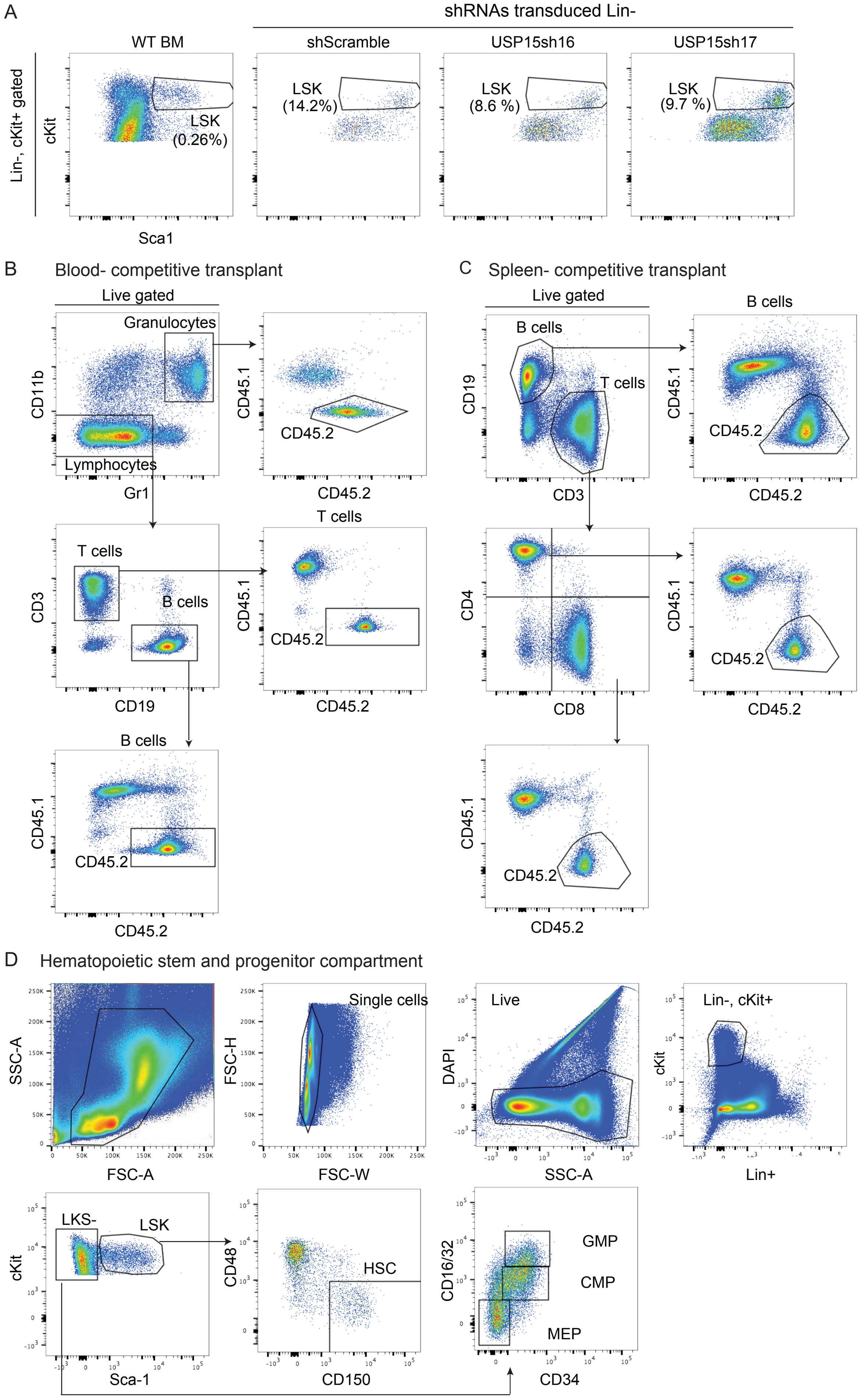

**Figure S3. Flow cytometry and gating strategies (relative to Figure 2C, 3, 4 nd 5).**

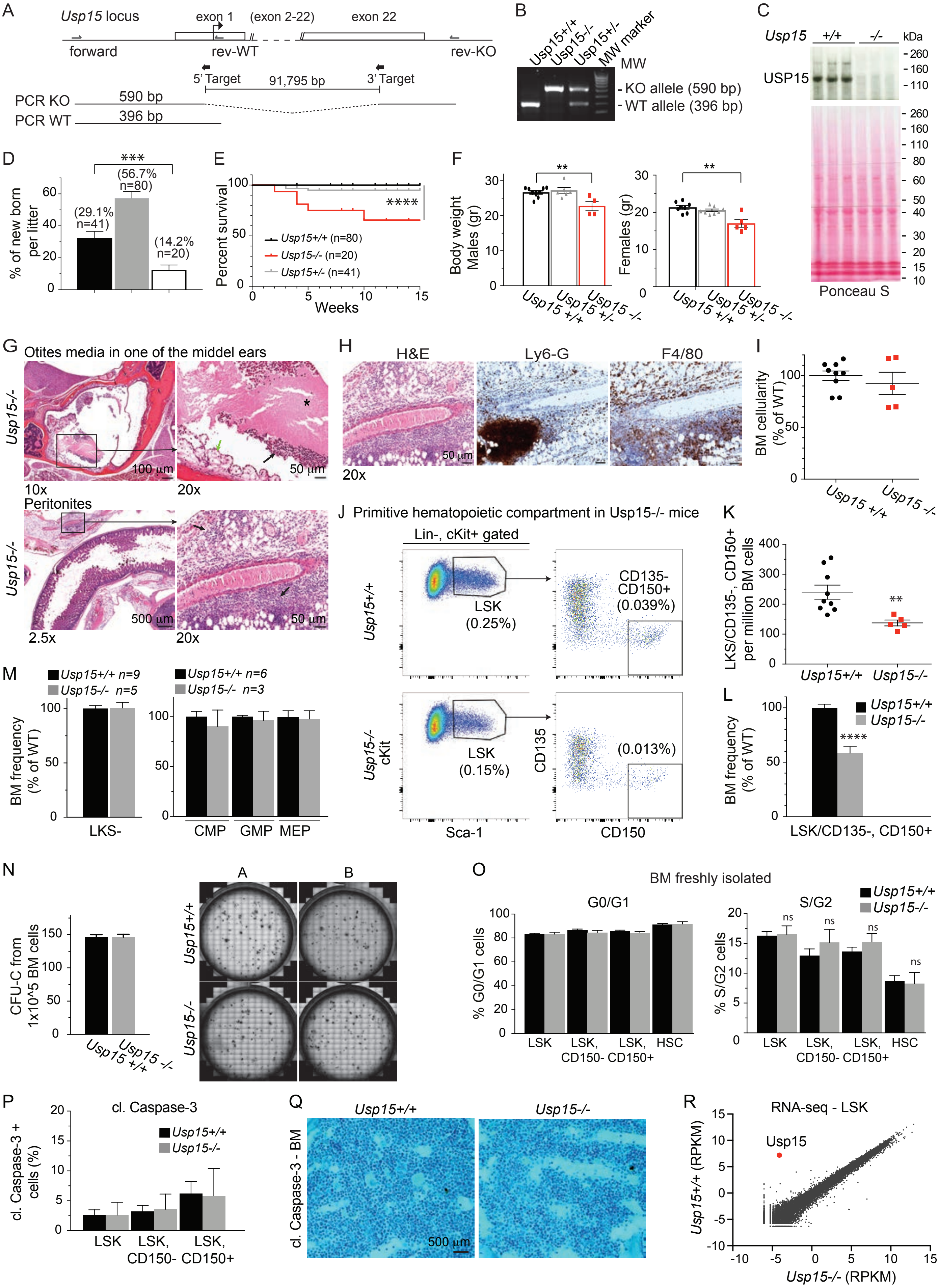

**Figure S4.Generation and initial characterization of Usp15 knockout mouse model (relative to Figure 4).**

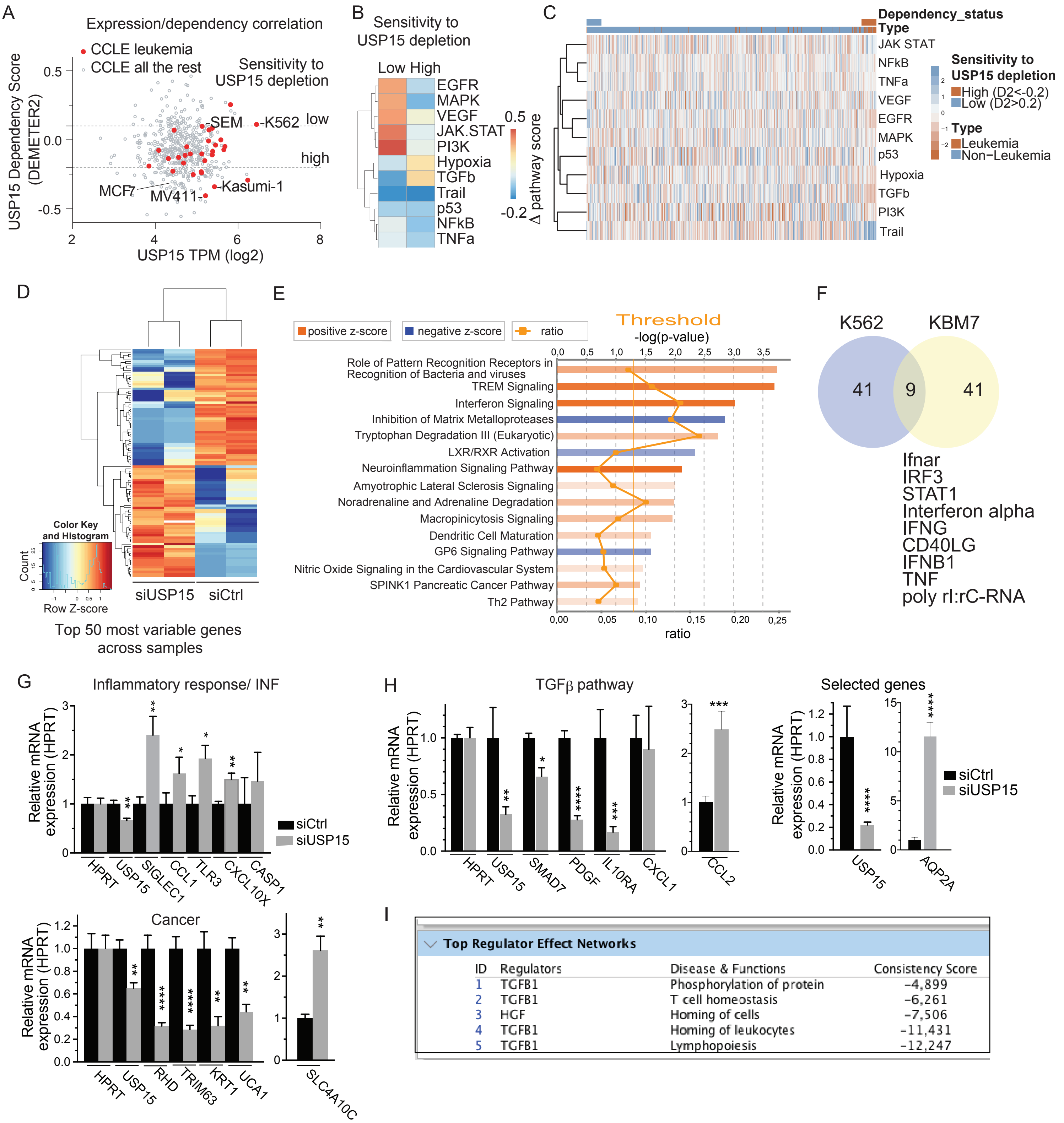

| ID | Regulators | Disease & Functions | Consistency Score |
|---|---|---|---|
| 1 | TGFB1 | Phosphorylation of protein | -4,899 |
| 2 | TGFB1 | T cell homeostasis | -6,261 |
| 3 | HGF | Homing of cells | -7,506 |
| 4 | TGFB1 | Homing of leukocytes | -11,431 |
| 5 | TGFB1 | Lymphopoiesis | -12,247 |

**Figure S5. USP15 RNAi in CML cell lines results in alteration of inflammatory signaling pathways.**

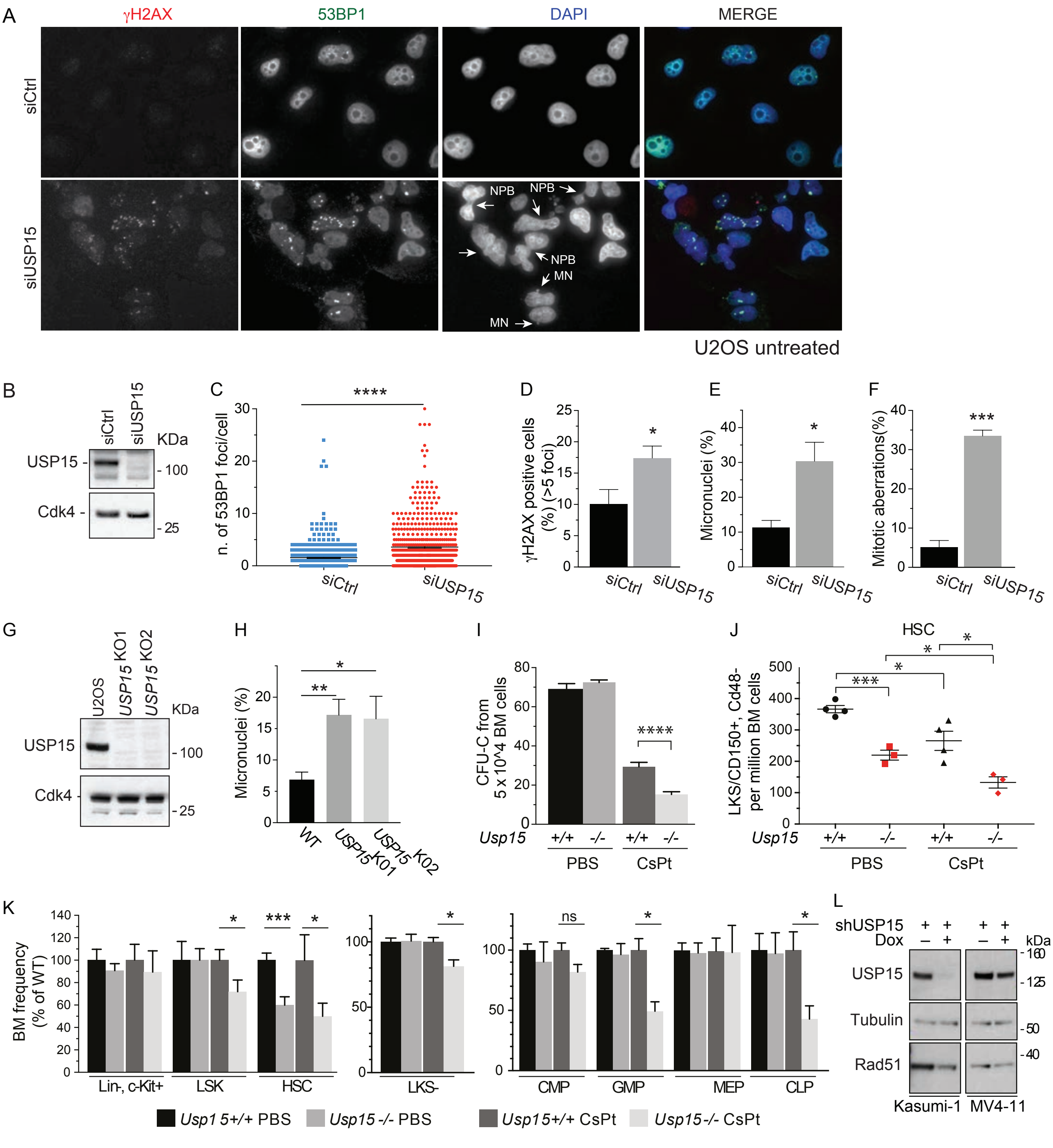

**Figure S6. USP15 loss enhances genotoxic stress *in vitro* in human cancer cells and *in vivo* in mouse hematopoietic stem and progenitor cells.**
